# DNA damage induced during mitosis undergoes DNA repair synthesis

**DOI:** 10.1101/2020.01.03.893784

**Authors:** Veronica Gomez Godinez, Sami Kabbara, Adria Sherman, Tao Wu, Shirli Cohen, Xiangduo Kong, Jose Luis Maravillas-Montero, Zhixia Shi, Daryl Preece, Kyoko Yokomori, Michael W. Berns

## Abstract

Understanding the mitotic DNA damage response (DDR) is critical to our comprehension of cancer, premature aging and developmental disorders which are marked by DNA repair deficiencies. In this study we use a micro-focused-laser to induce DNA damage in selected mitotic chromosomes to study the subsequent repair response. Our findings demonstrate that (1) mitotic cells are capable of DNA repair as evidenced by DNA synthesis at damage sites, (2) Repair is attenuated when DNA-PKcs and ATM are simultaneously compromised, (3) Laser damage may permit the observation of previously undetected DDR proteins when damage is elicited by other methods in mitosis, and (4) Twenty five percent of mitotic DNA-damaged cells undergo a subsequent mitosis. Together these findings suggest that mitotic DDR is more complex than previously thought and may involve factors from multiple repair pathways that are better understood in interphase.

## Introduction

DNA damage occurs naturally through various endogenous and exogenous processes. Unrepaired DNA can compromise genetic integrity leading to developmental disorders, cell death or cancer. Organisms have evolved a variety of pathways to respond to the damage. The vast majority of studies on DNA damage responses have been done during interphase of the cell cycle. However, understanding the DNA damage response (DDR) during mitosis is also important since mutations accumulated during mitosis can lead to chromosomal aberrations, genomic instability of daughter cells, senescence and eventual cell death [1–4].

Studies examining the extent of DDR activation and repair in mitosis have primarily assessed the cellular response to double strand breaks (DSBs). DSBs may be repaired by homologous recombination (HR) and non-homologous end joining (NHEJ). HR preserves genetic fidelity as it relies on a homologous template to restore the damaged DNA. On the other hand, NHEJ leads to ligation of broken ends which can lead to loss of genetic information. Studies examining the DDR of DSBs in mitosis found truncated DDR that does not lead to the accumulation of ubiquitin ligases as well as 53BP1 and BRCA1 at mitotic damage sites [2, 5–10]. Subsequent studies revealed that mitosis-specific phosphorylation of 53BP1 by polo-like kinase 1 (PLK1) block 53BP1 binding to chromatin [11, 12]. Furthermore, RAD51 and filament formation were found to be inhibited by CDK1 in mitosis [13, 14]. Taken together, DSB repair, both NHEJ and HR, were thought to be inhibited in mitosis.

Further, DNA synthesis has been investigated in early mitosis with respect to DNA damage resulting from replication stress. However, this form of repair has been shown to be dependent on a process that is only activated in very late G2/early prophase [15–22]. Cells treated with Aphidicolin in S phase had Rad52 and MUS81-EME1 dependent DNA synthesis at chromosome fragile sites during very early prophase [16, 17]. The MUS81-EME1 complex and BLM helicase are required for the restart of DNA synthesis after replication stress [18–20]. Interestingly, cells synchronized to prometaphase did not undergo replication stress induced DNA repair synthesis [16, 17]. Additionally, MUS81 is associated with chromosome fragile sites in prophase but at a decreased rate in metaphase [18]. This form of DNA synthesis has been termed MiDAS (Mitotic DNA repair synthesis) and should not be confused with the DNA repair synthesis described in this paper. The mechanism of replication stress induced DNA synthesis likely differs from that observed in response to damage elicited by other means and from damage induced in other phases such as prometaphase, metaphase and anaphase. Thus, the ability of mitotic cells to undergo DNA repair synthesis in response to damage elicited during mitosis remains to be elucidated.

The majority of DDR studies have utilized ionizing radiation or radiomimetic drugs to induce DNA damage and study the subsequent mechanisms of DNA repair. These methods of DNA damage induction result in genome-wide alterations that may lead to different DDRs. However, the laser has demonstrated to be very useful for DNA damage response studies because of its ability to target a submicron region within a specified chromosome region [23–29]. Interestingly, a laser micro-irradiation study conducted over forty years ago, showed that when the nucleolar organizer was specifically damaged in mitotic cells, a few of the irradiated cells were able to undergo a subsequent mitosis. Karyotype analysis revealed intact chromosomes with a deficiency of the nucleolar organizer [26, 30]. In today’s context, these results suggest that the cells most likely repaired mitotic DNA damage through NHEJ.

Studies by our lab and others have shown the ability of a diffraction-limited focused near-infrared (NIR) 780nm laser micro-beam to induce DSBs marked by γH2AX, phosphorylated Histone H2AX on Ser 139, and KU in both interphase and mitosis [29, 31–37]. In addition to γH2AX and KU at laser-damaged sites, we have demonstrated that ubiquitylation was also occurring at damage sites in mitosis [28, 29]. Though our findings differ from those that utilized ionizing radiation [5, 10], the laser micro-irradiation approach permits the visualization of proteins such as KU, that do not form ionization radiation induced foci (IRIF) [38]. Therefore, it is not surprising to see accumulation of DDR proteins at laser damaged regions in mitosis that have been previously thought to be excluded from mitotic DNA damage.

In the current study we systematically characterize the nature of mitotic DNA damage induced by the NIR laser and perform quantitative analysis of DNA repair in mitosis and subsequent G1 phase in human and rat kangaroo (*Potorous tridactylus*) cells. Under our conditions, the NIR laser micro-irradiation of mitotic chromosomes induces complex damage consisting of both strand breaks (DSBs and single-strand breaks (SSBs) and ultraviolet (UV)-crosslinking damage (pyrimidine dimers) similar to what was recently described for damage to interphase cells [39]. We demonstrate that factors from various repair pathways whose function is better understood in interphase are capable of responding to mitotic DNA damage. Our results also indicate that DSBs generated on metaphase chromosomes lead to clustering of various proteins involved in NHEJ and HR. We show that DNA repair of mitotic DNA damage is ongoing and persists into G1.

## Materials and Methods

### Reagents

See Supplemental S3 for a list of antibodies. Other reagents are listed under corresponding methods below.

### Cell Lines

Five human cell lines were utilized in this study. U-2 OS cells, referred to as U2OS in this paper, are an osteosarcoma cell line ATCC HTB 96 that was used for the majority of studies unless otherwise mentioned. CFPAC-1 ATCC CRL 1918, a line derived from cystic fibrosis pancreatic adenocarcinoma was also utilized. The isogenic cell lines, M059K ATCC 2365 and M059J ATCC 2366 were utilized to compare the mitotic DNA response when DNA-PKcs is absent (M059J). M059K contains the wild type form of DNA-PKcs making this line a good control for the DNA-PKcs mutant. Both cell lines come from a glioblastoma in the same patient. Rat kangaroo cells (PtK2) from the Potorous tridactylus were utilized due to their large chromosomes and strong adherence to the substrate that facilitate mitotic studies. These cells were grown in Advanced DMEM/F12 with 1% Glutamax and 10% Fetal Bovine Serum. All Human cells were grown in Advanced DMEM supplemented with 1% Glutamax and 10% Fetal Bovine Serum. All cells were maintained in a humidified 5% CO_2_ incubator. Cells were plated onto glass bottom gridded dishes from MatTek to a confluency of 40% and used 1-2 days post subculture. Experiments were carried out in medium containing 40ng/mL nocodazole. For inhibitory experiments the following concentrations were utilized: ATM inhibitor (10μM KU55933), DNA PKcs inhibitor (3μM NU7441), and PARP inhibitor (100μM Nu1025).

LASER Induced DNA Damage and Microscope Image Acquisition Mitotic chromosomes in live cells were irradiated using diffraction-limited (0.5 – 1 µm diameter) focal spots with a Coherent Mira 76 MHz 200 femtosecond micro- pulsed laser emitting at 780nm (Coherent Inc., Santa Clara CA). A series of beam expanders and mirrors coupled the beam into the right side port of a Zeiss Axiovert 200M inverted microscope. An X-Y fast scanning mirror was positioned in the beam path prior to entry into the microscope port to facilitate moving the focused laser beam toward the desired target. The beam was focused through a 63x (1.4 NA) Zeiss Plan-Apochromat oil objective. The irradiance of the laser was controlled through the use of a Glan-Thompson polarizer mounted on a motorized rotational stage. A Uniblitz mechanical shutter controlled the exposure time of the laser. A single chromosome within the cell was targeted by the laser unless otherwise noted. Chromosomes were exposed to the laser for a total of 10ms within the focal spot. This exposure resulted in 7.6x10^5^ pulses of light to a spot measuring 0.68μM in diameter at the selected irradiance of 2.8-3.2x10^11^W/cm^2^.

Images were collected using a Hamamatsu CCD Orca [33, 40, 41]. The polarizer, scanning mirror and shutter were controlled by software developed with LabView [42]. To determine the irradiance at the focal point, the transmission of the objective was measured using a modified dual objective method described previously [28]. The objective used in these experiments had a transmission of 0.50 at the laser wavelength used. Based upon the measured irradiance of 2.8- 3.2x10^11^W/cm^2^, the damage mechanism is likely of a multiphoton nature, either 2- photon or 3-photon, or a combination of both.

### Immunostaining

Cells were fixed with 4% paraformaldehyde in phosphate buffered saline for 20 minutes. Time to fixation after laser exposure varied according to the experiment. Cells were permeabilized overnight with blocking buffer containing 0.1% TritonX and 5% fetal bovine serum in phosphate buffered saline followed by staining with primary antibodies. Supplemental S3 shows a list of antibodies used. For most primary antibodies a 1:500 dilution was applied. Secondary antibodies against primaries were: Alexa-488 goat anti-mouse (Invitrogen, Carlsbad, CA), and Cy3 goat anti-rabbit (Invitrogen, Carlsbad, CA) at dilutions of ½000.

### DNA synthesis detection

To test for repair/DNA synthesis, cells were incubated with 10µM EdU (5-ethynyl- 2’-deoxyuridine) 1-20min before laser exposure. A 10mM EdU stock was prepared according to protocol (Invitrogen catalogue #C10339). Cells that require prolonged mitosis were concurrently incubated with colcemid and EdU between 1-20 minutes prior to irradiation. Cells were fixed at time points ranging from 10-120 minutes after irradiation with 4% paraformaldehyde in PBS for 5-10minutes, and followed by blocking buffer containing 10% fetal bovine serum and 0.2% Saponin in PBS for 30minutes

### Terminal deoxynucleotidyl transferase (TdT) dUTP Nick-End Labeling (TUNEL) assay

DNA end-breaks were detected at sites damaged by the laser in mitotic chromosomes by TUNEL assay; dUTP labeling of exposed 3’-ends of DNA strands. The assay was followed according to the manufacturers protocol (Roche Applied Science).

### Image Analysis

Tiff images were analyzed and subsequently edited to enhance the contrast and intensity using Image J software [43]. Mean pixel intensities(MPI) for laser DNA damaged regions were measured prior to contrast enhancement. The background was identified as the region outside of the DNA damage area and the mean pixel intensity of this area was subtracted from the fluorescence intensity of the lines or spots containing DNA damage in order to calculate the average pixel intensity at the damaged region. Positive signal for fluorescent markers was based on mean pixel values being higher than the level of background at undamaged chromosomes.

### UV Induced DNA damage

U2OS cells were subjected to 254 nm light from a UVG-11 Compact UV Lamp by placing the lamp directly over 50mm cell dishes for 20 seconds. The power of the lamp was monitored with a PM100 ThorLabs Power Meter equipped with a S120UV sensor. The average power was measured before each experiment and determined to be 6mW. Prior to UV lamp exposure cells were switched into phenol red free Hanks buffered saline (Invitrogen). After UV exposure cells were either immediately fixed with 4% Paraformaldehyde or placed into a 37 C incubator prior to fixation.

### Pyrimidine dimer quantification

Mitotic U2OS were collected via mitotic shake off after synchronization with 9uM CDK1 inhibitor (Calbiochem; RO-3306). CDK1 inhibition results in arrest at G2; upon removal of inhibitor cells entered mitosis. Collected cells were exposed to 265nm UV light from a UV lamp UVG-11 (Science Company) for 20s in phenol red free medium. Cells were then separated into three aliquots and lysed at 30, 60 and 90 minutes after UV exposure. DNAzol Reagent (Life Technologies) was added to each sample for lysis and DNA isolation according to the manufacturer’s protocol. Isolated DNA samples were quantified using a Nanodrop 2000c (Thermo Scientific) and diluted to a working concentration of 2.0ug/ml in cold PBS. DNA samples were further diluted to plate on 96-well DNA High-Binding Plates. An OxiSelect UV-Induced DNA Damage ELISA Combo Kit was utilized to determine the concentration of pyrimidine dimers in each sample well (Cell Biolabs Inc). Plates were read with a Biotek Conquer ELX800 plate reader (Biotek Inc).

### Statistical Analysis

Prism 7 for MAC OS was utilized for all statistical analysis. A T test was performed for comparisons to controls unless otherwise noted. Values were considered significant if P <0.05.

## Results

### The laser induces complex DNA damage on mitotic chromosomes

Optimal laser parameters for the detection and consistent production of DNA damage in mitotic chromosomes were obtained by (a) varying the irradiance and comparing phase contrast image changes, and, (b) assaying for Nbs1 accumulation and γH2AX production. For this study we utilized irradiances of 2.8- 3.2x10^11^W/cm^2^ unless otherwise noted. These irradiances allowed us to immediately determine that chromosomes were effectively damaged due to rapid phase contrast changes in the irradiated chromosome regions (Fig 1A arrows at 4 and 6s). Dark material, which may reflect phase separation is visible at 16s post laser exposure. In a different example, of a cell fixed ∼5s after the laser, we see that γH2AX surrounds an area targeted by the laser (Fig 1B). In a previous study we showed that dark material is a result of the accumulation of DDR and that γH2AX may surround the DDR and or overlap with them [34]. Additionally, at these irradiances γH2AX and Nbs1 were detected nearly 100% of the time (48 of 48 cells) and (16 of 17 cells) respectively.

**Fig 1.**
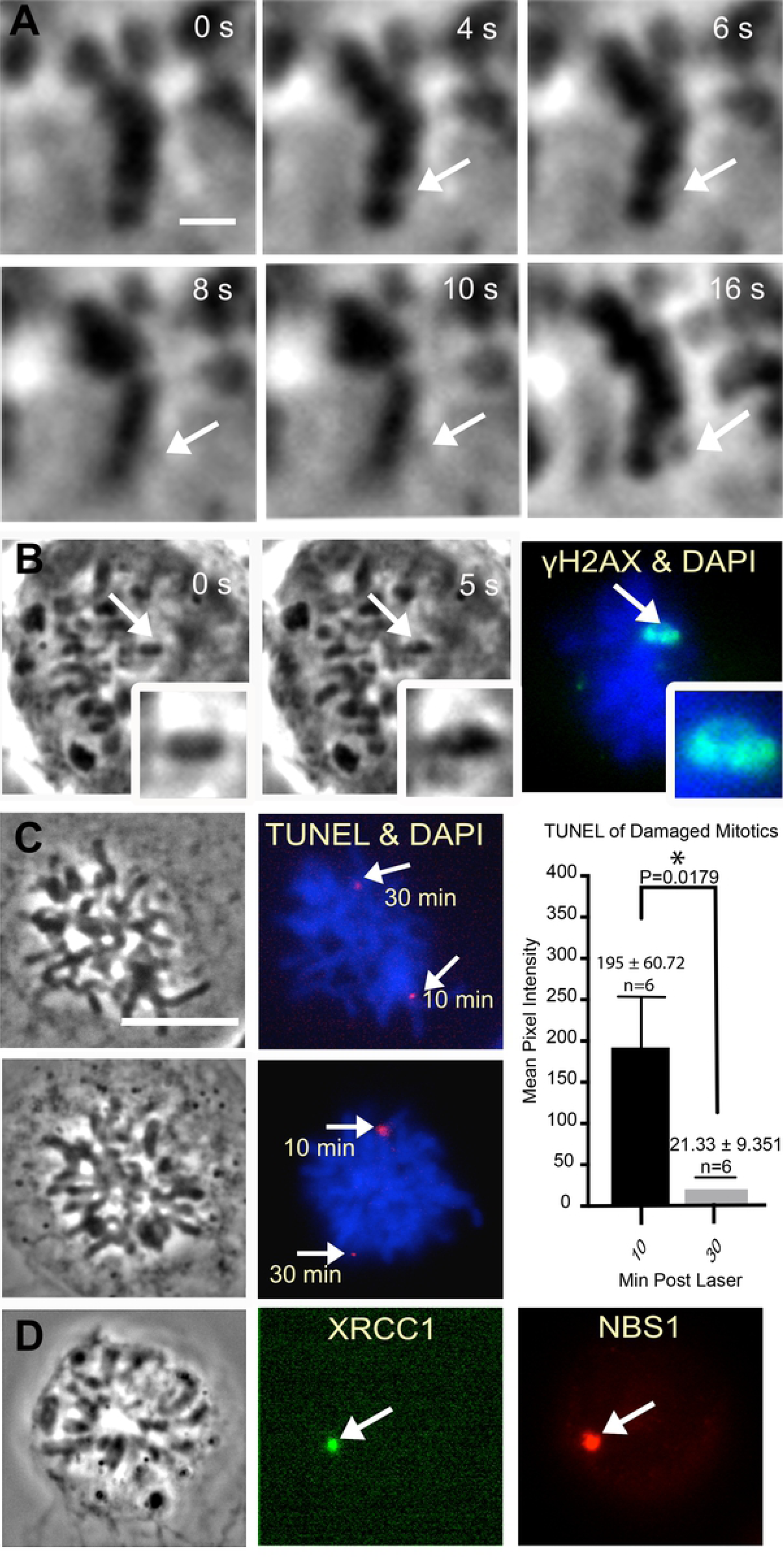
Characterization of Laser induced DNA damage. (A) At the selected irradiance of 3.0x1011W/cm2 phase contrast changes are observed at a laser damage site on prometaphase chromosomes in nocodazole synchronized U2OS cells (4 and 6s). At 16s post laser (see arrows) phase dark material is apparent. Scale bar=1μm. (B) A metaphase cell that was fixed ∼5s after the laser and stained for γH2AX (arrows and inset). This cell was fixed prior to dark material accumulation. Paling is observed in the inset. γH2AX is formed around the region damaged by the laser. (C) Positive TUNEL is seen at laser damaged regions (arrows) in prometaphase cells incubated with nocodazole. Two different chromosomes were damaged at different time points within the same cell. The first damage was induced 30 minutes prior to fixation and the second damage was induced 20 minutes after the first damage and fixed after 10 minutes. The TUNEL signal at the second damage site was brighter than the signal at the first damage. A graph of TUNEL signal on the right is the average of six cells per category. Scale bar=1μm. (D) XRCC1 and NBS1 co-localize to chromosome damage in a cell fixed 25 minutes post laser.

The type of DNA damage induced by the laser on mitotic cells was assessed by immuno-staining for (1) DNA damage response proteins, (2) damaged bases, and (3) the TUNEL assay. Experiments were carried out in human U2OS cells unless otherwise mentioned. Positive TUNEL at laser induced DNA damage sites demonstrated the presence of END breaks Fig 1C, (magenta). TUNEL signal was higher in chromosomes fixed 10 minutes post laser when compared to those fixed 30 minutes post laser. These results indicate that factors may have bound broken ends.

Several DDR proteins were found at damaged chromosomes which provide information on the type of damage created by the laser. KU heterodimer confirmed the presence of double strand breaks (DSBs) (Figs 2D, 4A and B, and S1). Pyrimidine dimers (CPD) as a result of the laser exposure were also detected and will be discussed further in this paper. XRCC1, which is involved in single strand break (SSB) repair, nucleotide excision repair (NER) and base excision repair (BER) localized to laser-damaged DNA (Fig 1D)[44]. However, we failed to detect significant base damage at laser targeted regions at the irradiance range of 2.8- 3.2 x10^11^ W/cm^2^ using an antibody specific for 8-Oxoguanine (8-oxoG) (Trevigen) (S1 A). Nevertheless, a key component of BER, APE-1 was detected at laser damage sites (S1 C).

**Fig 2.**
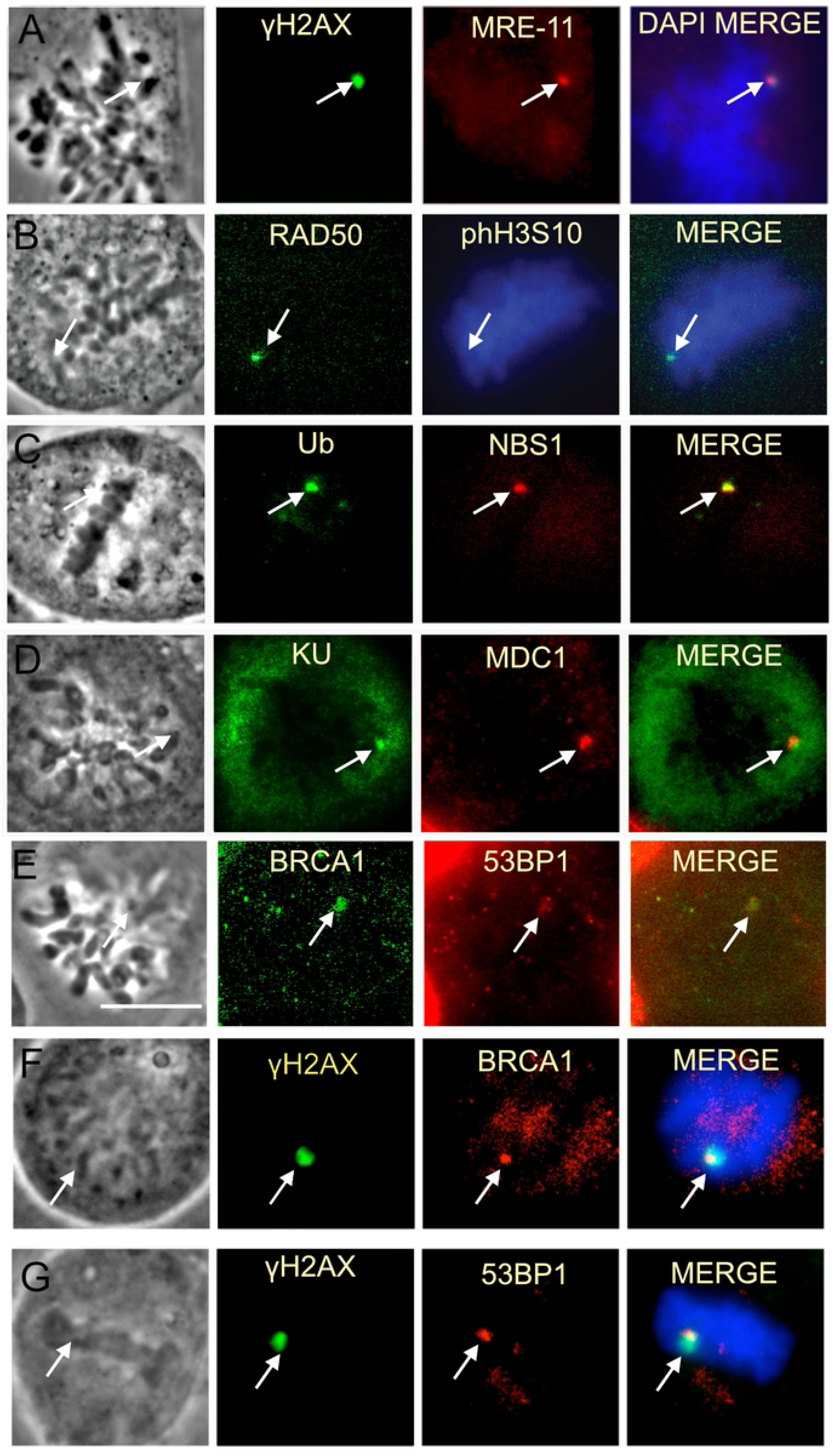
Mitotic laser induced DNA damage leads to the recruitment of various DDR proteins including some not previously observed on mitotic DNA damage. (A-C) The MRN complex (MRE-11, Rad50, Nbs1) forms at laser damaged chromosomes. Nocodazole synchronized U2OS cells are shown. MRE- 11(n=4), Rad50(n=3), NBS1(n=16). (C-E) Ub(n=19), KU(n=23), MDC1(n=7), 53BP1(n=6) and BRCA1(n=10) were also observed at chromosomes damaged by the laser. Scale bar=10μ m. (E and F) BRCA1 immunostaining using two different antibodies. (E) BRCA1 ab16780 antibody from Abcam. (F) BRCA1 OP107 antibody from Calbiochem. (F-G) show the localization of BRCA1 and 53bp1 with respect to yH2AX.

### Mitotic cells undergo DNA Repair synthesis

Since DNA repair generally has been perceived to be inhibited in mitosis, we directly monitored the repair event in mitotic cells by the incorporation of the thymidine analogue 5-ethynyl-2’-deoxyuridine (EdU). For these experiments, cells were incubated either 10 minutes prior to laser or 10 minutes post laser. For both conditions cells were fixed 30 minutes post laser (Fig 3A). Cells damaged in metaphase were capable of incorporating the analogues, and this incorporation was surrounded by γH2AX known to spread to the neighboring chromatin (Fig 3B)[45, 46]. Cells incubated with EdU prior to the laser showed significant repair during the first 10 minutes post laser (Fig 3C). We found that DNA repair synthesis is not restricted to the initial 10 minutes post laser as cells incubated with EdU post laser were still positive for EdU albeit weaker. demonstrating ongoing DNA synthesis. The incorporation of the EdU (i.e. repair) was observed in multiple cell lines during mitosis: U2OS (Fig 3B and C), and the Isogenic cell lines M059K & M059J (Fig 3D top panel; 3D bottom panel, respectively).

**Fig 3.**
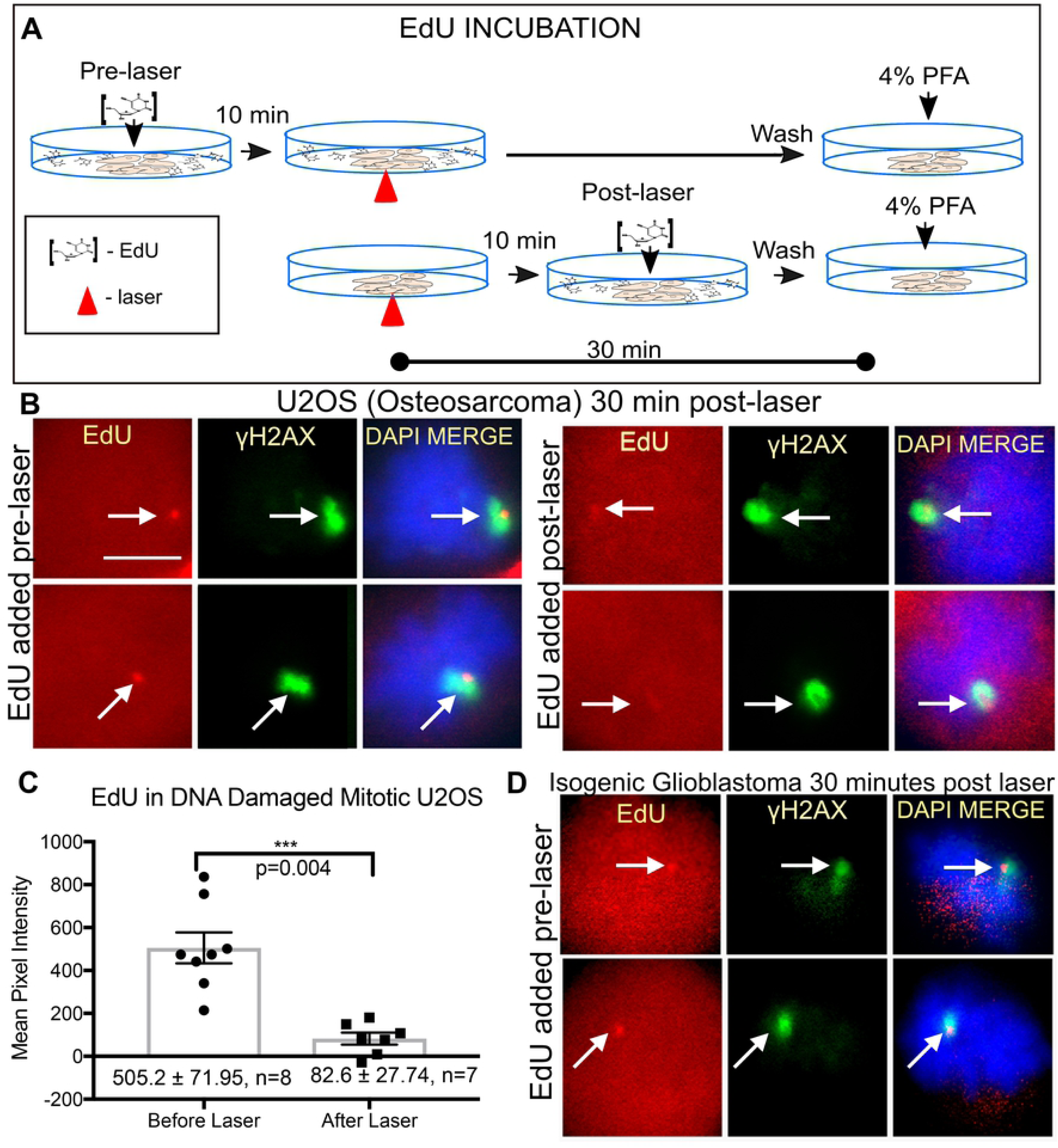
Mitotic cells undergo DNA synthesis repair. (A) Schematic of EdU incubation for experimental results shown in (B-D). In the first scenario EdU was added to cells 10 minutes prior to the laser to allow penetration into cells before damage. In the second scenario EdU was added to cells 10 minutes post laser to test whether repair is ongoing. All cells were fixed 30 minutes post laser. (B) Immunofluorescence images of nocodazole synchronized U2OS (Osteosarcoma) cells damaged under the conditions depicted in the schematic above were stained for EdU and γH2AX. γH2AX partially overlaps and surrounds EdU. Scale bar= 10μm (C) Quantifications of EdU intensity at the damage site in cells incubated pre or post laser. (D) Isogenic Glioblastoma cells M059K(top panel) and M059J(bottom panel) were stained for EdU and γH2AX.

### Complete assembly of Non-homologous End Joining factors in response to mitotic DNA damage

In an effort to identify what components of the DNA damage response may be active following laser damage to mitotic chromosomes, several damage sensors, adaptor proteins and transducers were tested for their ability to cluster to laser damage sites. Our evaluation begins with DSB repair pathway proteins. DSBs are amongst the most deleterious in that they can result in chromosomal translocations if left unrepaired [47].

During interphase the Mre11-Rad50-Nbs1 (MRN) complex is one of the first factors to recognize DSBs [48, 49]. At the selected irradiance all three components of the MRN complex are detected at laser-induced damage sites in mitotic cells (Fig 2A-C). Although, BRCA1, 53BP1 and Ubiquitin (Ub) responses have been observed to be attenuated in mitosis, [50, 51] we previously showed an Ub signal at NIR laser-induced damage sites in PtK1 cells [28]. In the present study, Ub accumulation was also observed in mitotic U2OS cells (Fig 2C). Ub response at damage sites is critical for localization of BRCA1 and 53BP1 at DSBs in interphase nuclei [52, 53]. Correlating with the presence of the Ub signal at mitotic damage sites, we observed the accumulation of BRCA1 and 53BP1 at mitotic damage sites (Fig 2E-G). Our results indicate that Ub, BRCA1 and 53BP1 can be recruited to highly clustered laser-induced damage sites on mitotic chromosomes.

DSBs may also be repaired by non-homologous end joining (NHEJ) which is an error-prone DSB repair pathway that leads to ligation of broken ends. NHEJ initiation has been observed in mitotic DNA lesions using a NIR laser in two separate studies [28, 38]. In our studies KU accumulated during the first 5-15 min post laser (Fig 4A & B). In contrast, the recruitment of DNA ligase IV that mediates end joining at a later step of NHEJ was not apparent until 20 minutes post irradiation (Fig 4A and B). DNA PKcs and XRCC4, the binding partner of ligase IV, was also detected at laser damage (Fig 4C and D). Therefore complete assembly of NHEJ factors may occur in mitosis.

**Fig 4.**
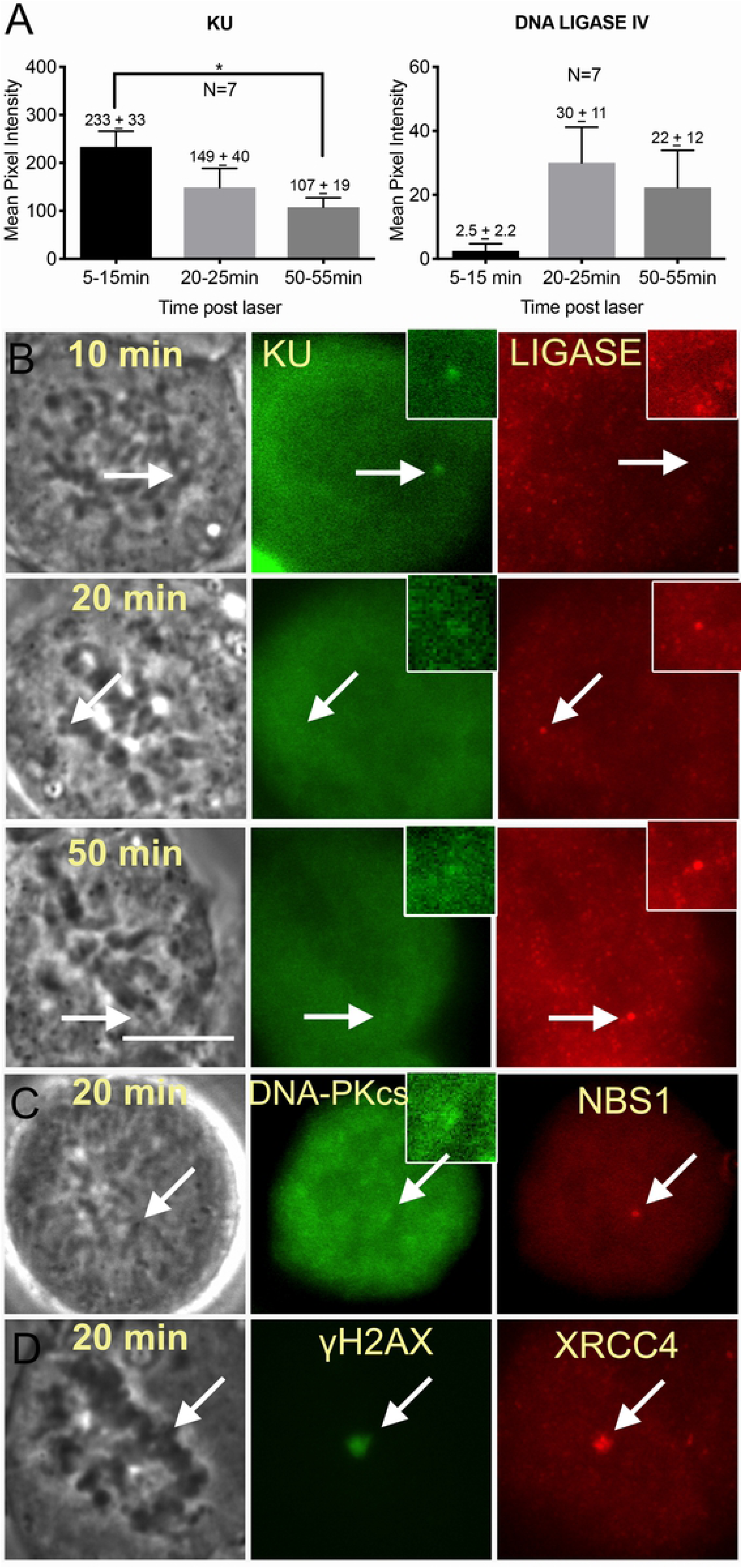
NHEJ factors cluster at Mitotic DNA damage. (A) KU and LIGASE signal intensity quantified in U2OS at various times post laser. Cells were maintained in nocodazole throughout experiments. N=7 for cells fixed 5-15 and 20-25 minutes post laser, N=9 for cells fixed 50-55 minutes post laser (B) Cells stained for KU and LIGASE IV. An arrow depicts the area targeted by the laser and a region that is magnified as an inset. Scale bar=10 μm. (C) DNA-Pkcs clusters to damaged U2OS chromosomes. An inset depicts a magnified view of DNA-PKcs at the cut site. (D) XRCC4 and γH2AX at laser damaged region.

Classic NHEJ is dependent on DNA-PKcs. We investigated the contribution of NHEJ on DNA synthesis repair by utilizing DNA-PKcs deficient (M059J) and isogenic DNA-PKcs-positive (MO59K) cell lines. Immunostaining of DNA-Pkcs in the isogenic lines confirmed its presence in M059K and absence in M059J (S2 A). Nevertheless, DNA repair synthesis was observed in both cell lines (Fig 2D and 5B). Similarly, U2OS cells treated with 3 μM DNA-PKcs inhibitor NU 7441 showed no significant difference when compared to control cells (Fig 5A DMSO vs DNA- PKcs inhibitor).

**Fig 5.**
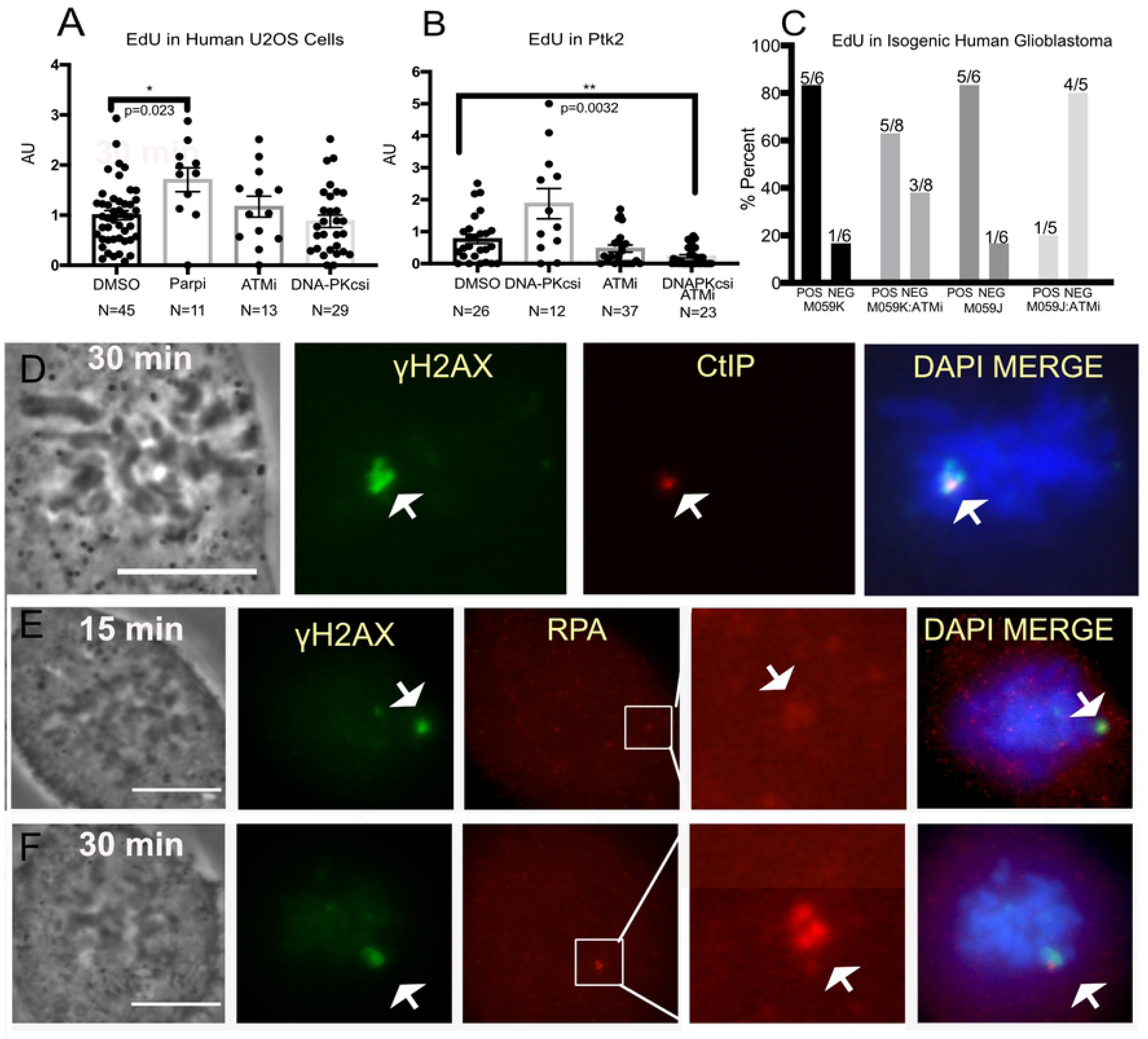
Mitotic DNA synthesis on DNA damaged chromosomes of cells with compromised ATM, DNA PKcs or PARP activity. (A) Nocodazole synchronized U2OS cells treated with inhibitors for PARP (100μM NU1025), ATM (10μM Ku55933) and DNA PKcs (3μM NU7741)were assessed for DNA synthesis. Each experiment was normalized to the mean of the corresponding DMSO control. (B) Synchronized PtK2 cells treated with DMSO, ATM inhibitor, and combined DNA- PKcs and ATM. (C) Nocodazole synchronized human isogenic cell lines M059K and M0959J (DNA-PKcs deficient) were treated with ATM inhibitor. The percent of positive cells is plotted. Above each bar is the number of cells per category. (D) CtIP and (E) RPA were found on laser-damaged chromosome regions on U2OS cells fixed 15(top panel) and 30(bottom panel) minutes post laser.

Alternative-NHEJ (alt-NHEJ) may also repair DSBs when DNA-PKcs is compromised [54]. Poly (ADP-ribose) polymerase 1 (PARP1) has been shown to play a pivotal role in alt-NHEJ [55, 56]. Therefore we inhibited PARP in U2OS cells with 100μM NU 1025 and tested for effective inhibition by immunostaining for poly(ADP-ribose) (S2 B). S2 D depicts an untreated U2OS mitotic cell positive with PARP staining at the damage site. Interestingly EdU results from cells synchronized with nocodazole suggest that PARP inhibition may lead to greater DNA synthesis (Fig 5A, DMSO vs PARPi). Since PARP inhibition is known to stimulate c-NHEJ, the effect of PARPi on EdU incorporation was examined in DNA-PKcs deficient cells [39, 57, 58]. PARP inhibition did not abolish DNA synthesis at mitotic DNA damage sites (S2 C). These results reveal that PARP signaling plays a role in suppression of DNA repair during mitosis.

### Homologous Recombination is activated in mitosis and may lead to initiation of RAD51 filament formation in the absence of functional DNA-PKcs or CDK1

Homologous recombination may preserve genomic integrity during the repair of DSBs. This process relies on resection of the damaged DNA ends and a homologous template to synthesize new DNA and preserve genomic integrity. ATM kinase is key to the activation of this repair pathway [59, 60]. ATM inhibition by 10μM KU 55933 in M059J cells significantly attenuated DNA synthesis at mitotic damage sites (Fig 5B MO59J:ATMi). Similar results were obtained in mitotic PtK2 cells treated with both DNA-Pkcs and ATM inhibitors (Fig 5C). Since ATM was shown to stimulate HR repair in interphase [59, 60], these results raise the possibility that when both c-and alt-NHEJ pathways are inhibited, HR factors may contribute to DNA repair during mitosis.

A key first step towards HR repair involves DNA end resection through CtIP followed by RPA binding to the resected DNA ends [61]. Consistent with previous studies that used meiotic Xenopus extract to monitor DSB repair, we observed CtIP and RPA at mitotic laser damage sites suggesting ongoing end resection (Fig 5 D and E) [62].

Downstream of end resection and RPA binding is RAD51 filament formation for homologous strand invasion. Previously, RAD51 was reported to not accumulate to damaged meiotic chromatin from X. laevis egg extract unless CDK1 was inhibited [62]. Similarly, we did not observe RAD51 filament formation in U2OS mitotic cells synchronized with nocodazole (Fig 6A and C). In contrast, RAD51 accumulation was observed at the laser damage sites in interphase cells fixed at 25 minutes post laser damage. Similar results were obtained with a different human cell line, CFPAC-1, indicating that this is not a cell type-specific phenomenon (S2 E and F).

**Fig 6.**
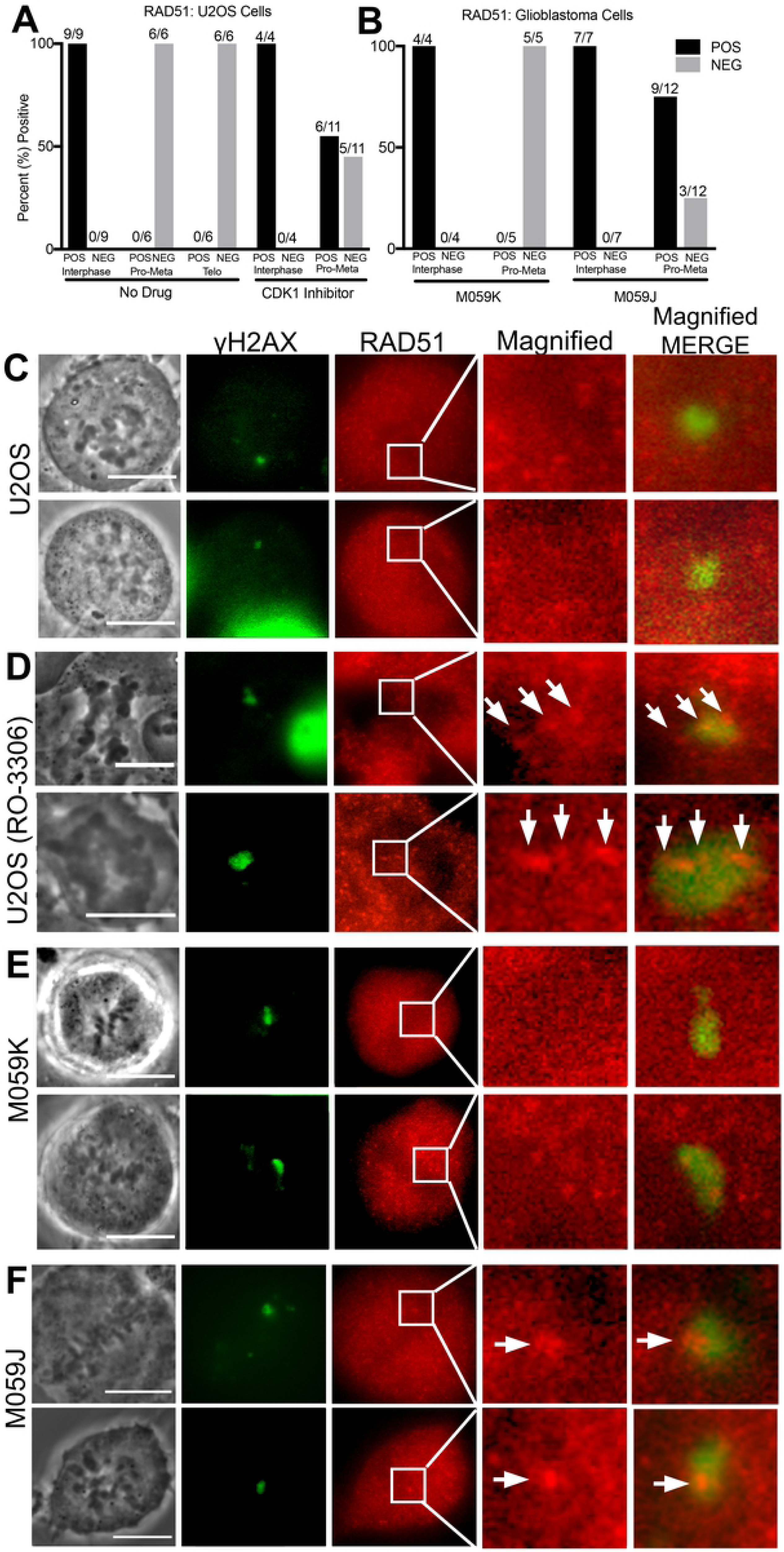
DNA damaged chromosome regions are devoid of RAD51. (A) Percent of U2OS cells positive for RAD51 according to mitotic phase. CDK1 inhibition causes some cells to present mean pixel values above background, MPI=48 ± 41in six out of eleven cells. (B) DNA Pkcs deficient cells, (M059J) also show a greater likelihood of RAD51 above background levels when compared to U2OS. Nine out of twelve had positive RAD51, MPI=115 ± 69. (C) U2OS mitotic cells stained for γh2AX and RAD51. (D) U2OS treated with CDK1 inhibitor underwent premature cytokinesis and chromosome de-condensation. Nevertheless, RAD51 co-localized with γh2AX and appears dotted along a track in an enlarged image of a boxed region shown on the RAD51 column. (E) The isogenic glioblastoma lines M059K and (F) M059J were DNA damaged with the laser and stained for γh2AX and RAD51. (F) M059J cells (deficient in DNA-PKcs) show RAD51 at the damage spot. Scale bar=10μ m

CDK1 was shown to play a role in HR inhibition in mitosis [9, 11, 62, 63]. Cells treated with 10μM CDK1 inhibitor R0-3306 underwent premature cytokinesis and or chromosome de-condensation within 5 to 10 minutes of inhibitor addition (Fig 6D). A proportion of CDK1 inhibited mitotic U2OS cells demonstrated slightly higher fluorescence pixel values above background (48 ± 41) than control cells whose values were negative (Fig 6A and D). RAD51 appears filamentous at damage sites showing positive RAD51 staining (Fig 6D, arrows on magnified view). However, the levels of fluorescence intensity were not in the same positive range of RAD51 (1409 ± 660 mean pixel intensity) seen in interphase cells. Thus, it would appear that other pathways independent of CDK1 activity may regulate RAD51 suppression.

Interestingly, in the absence of DNA-PKcs, most cells (9 of 12) showed some RAD51 at damage sites (MPI=115 ± 69) with a large standard deviation and a range of 76 to 1288 MPI (MPI=680). These results suggest that mitotic DNA-PKcs also regulates RAD51 accumulation at clustered damage sites (Fig 6B and F). Interestingly the staining pattern of RAD51 differs from that observed in CDK1 inhibited U2OS cells (compare magnified view i.e. 10x of Fig 6D and F). M059K cells did not show RAD51 that localized to the damage area (Fig 6E).

Taken together, in contrast to NHEJ factor assembly, only the early part of the HR pathway proteins accumulate to damaged DNA on mitotic chromosomes. CtiP and RPA both increase over time suggesting the formation of more strand breaks or end resection. RAD51 binding may be stimulated in mitosis when DNA-PKcs or CDK1 are compromised. Nevertheless, Rad21 was not observed at damaged regions (S2 G). This is possibly due to the fact that cohesin is destabilized during mitosis which only promotes HR between sister chromatids but not other types of HR[64, 65]

### Mitotic DNA repair synthesis is not a laser specific phenomenon

Our EdU labeling results strongly indicate that there is repair of laser-induced damage in mitosis. To confirm that mitotic DNA synthesis repair can occur in cells damaged by other means we exposed cells to UV light from a lamp and then isolated by FACS using an antibody specific for phosphorylated histone H3 Serine 10 (phospho-H3S10) (Fig 7A red). Phospho-H3S10 is greatest during mitosis and therefore the mitotic population can be easily separated from the interphase population. Mitotic cells were plotted against EdU fluorescence intensity (Fig 7A, bottom panels). The scatter plots reveal that 50 percent of cells stained positive for EdU in response to UV when compared to 30 percent of control cells. A histogram of the same results in Fig 7B depict the increase in proportion of cells staining positive for EdU has inceased (red histogram compared to blue). The results indicate that UV damage repair can occur in mitotic cells and mitotic repair is not a laser damage specific phenomenon. Amongst our findings are results showing that mitotic cells are capable of removing pyrimidine dimers (CPD) as confirmed through ELISA of UV damaged cells (Fig 7C). Similarly, cells damaged by the laser demonstrated a decrease of pyrimidine dimers (Fig 7D). PtK2 and U2OS cells stained for CPD are shown in Fig 7E and F, respectively.

**Fig 7.**
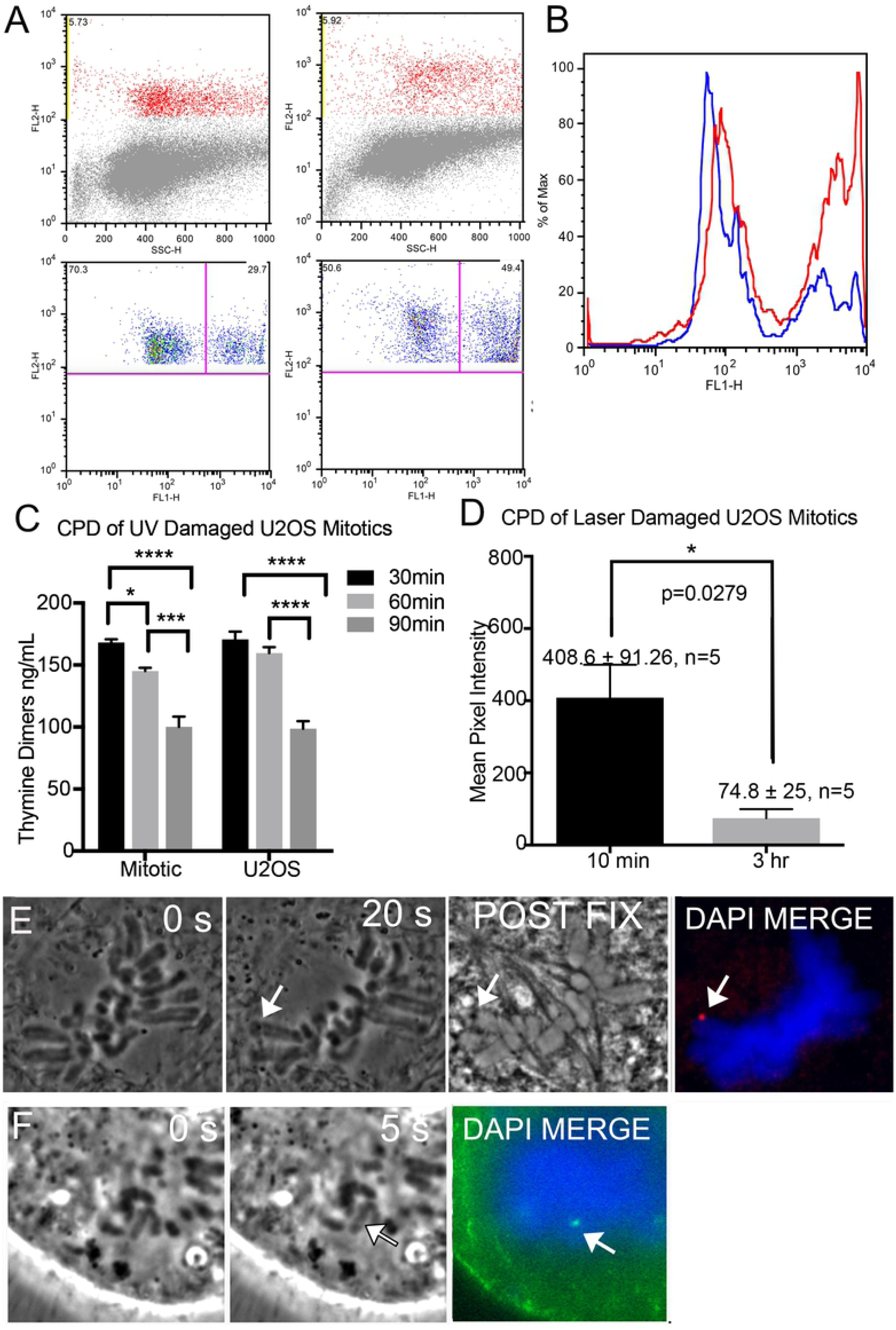
UV induced DNA damage repair in mitosis. (A) FACS of U2OS cells stained for mitotic marker, phospho-H3S10 (y-axis) plotted against side scatter (x-axis) on the top two panels. Cells that stained positive for phospho-H3S10 were plotted against EdU (x-axis) in the bottom panels. The right quadrant of each plot shows mitotic cells that stained positive for EdU. A greater proportion of cells are positive for EdU following UV exposure. Compare 30% without UV to 50% with UV. (B) A histogram of both populations, of cells, damaged/UV exposed in red and undamaged in blue to show the way the populations shift towards greater EdU signal after UV exposure. (C) ELISA of a population of non-laser UV exposed synchronized mitotic and interphase cells collected at 30, 60, and 90 minutes post exposure. N=3 replicates for mitotic populations and N=2 for interphase cells. (D) Quantification of CPD intensity in U2OS mitotic cells compared at 10 minutes and 3hours post laser, N=5 per category. (E) A mitotic PtK2 whose chromosome was damaged by the laser(arrow). A post fixation phase image of the cell shows a dark spot at the laser cut site. Cyclo-butane pyrimidine dimers (red) are seen at the exposure site. (F) A U2OS chromosome positive for cyclo-butane pyrimidine dimers (green) at the damaged site.

### Mitotic DNA damage is carried into interphase

The ability of clustered mitotic DNA damage to accumulate RAD51 post mitosis was investigated. RAD51 was not detectable in our studies unless CDK1 and or DNA-PKcs activity was compromised. Therefore, cells were examined for the ability of HR factors to recruit to laser damage created in mitosis once the cells had entered G1. We found that in the following G1 phase, RAD51 does accumulate to the DNA damage produced in the preceding mitosis (Fig 8A and B). EdU colocalized with RAD51 indicating the possibility that HR may be responsible for some of the incorporation and that repair is ongoing (Fig 8B and 8D). A cell damaged in metaphase and fixed 40 hours post mitosis is still undergoing DNA repair synthesis(Fig 8D).

**Fig 8.**
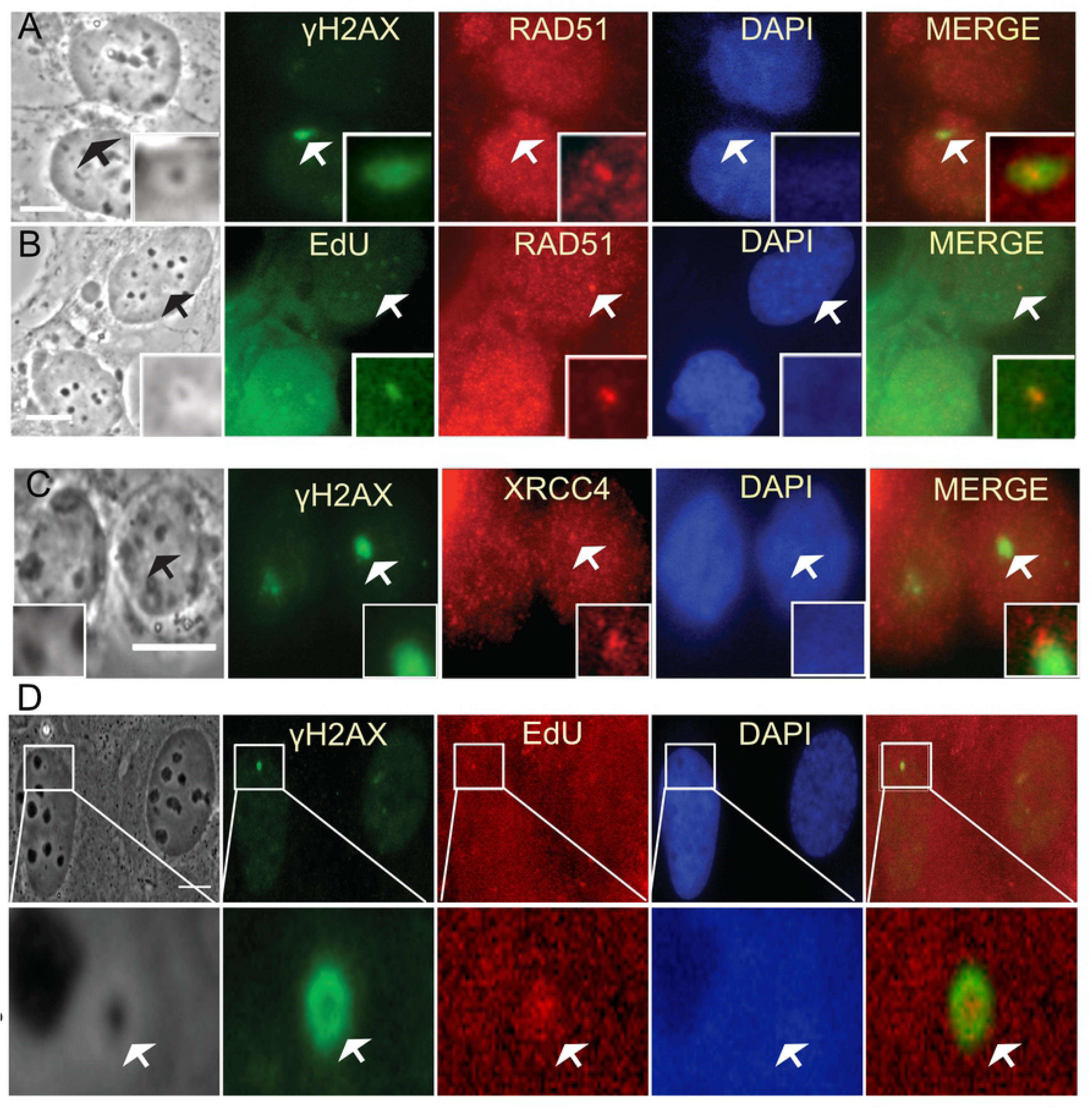
Mitotic DNA damage carried into G1 suggests ongoing repair. Cells damaged in mitosis where fixed 2 hours post division and stained for downstream HR and NHEJ factors as well as for EdU. (A) RAD51 and γ H2AX localize to the same area of a G1 cell. (B) EdU and RAD51 co-localize with each other and a phase dark spot of a G1 cell. (C) XRCC4 and γ H2AX slightly overlap in a G1 cell. Insets depict a magnified view of damage spots pointed out in arrows. (D) Forty hours post mitosis a cell has a phase dark spot that is surrounded by γ H2AX and that co-localizes with EdU. Scale bar= 10μm.

Additionally, we assessed whether NHEJ factors are still present in G1 from damage created in mitosis. Immuno-staining for XRCC4 showed that, in fact, XRCC4 is still present at G1 (Fig 8C). Thus, it seems that the cell may be trying to repair clustered laser damage utilizing factors from NHEJ and HR. Previously we reported that BRCA1 and 53BP1 were also observed post mitosis[28]. This result further supports repair is ongoing.

We investigated the ability of cells damaged in metaphase, anaphase and G1 to complete mitosis and enter a subsequent mitosis. For this, undamaged U2OS cells followed under our culture conditions showed an average division time of 37 ± 5 hours post cytokinesis. Values were calculated by taking the time of cytokinesis and following a cell until its entry into the subsequent mitosis (S3 A controls, N=11 cells). As a result, damaged cells were followed for a minimum of 40 hrs.

The majority of mitotic cells damaged with the laser entered G1, 26 of 28 of cells damaged in metaphase and 8 of 10 cells damaged in anaphase (Fig 9A and B). One cell damaged in metaphase died. The daughters of cells that divided were followed and entry into mitosis i.e. completion of a cell cycle was assessed within an observation window up to 40-47 hours post DNA damage. A proportion (21%) of daughter cells with damage inflicted in pro-metaphase underwent a subsequent mitosis, compared to 37% for daughter cells carrying damage elicited in anaphase. Examples of time-lapse analysis of cells damaged in metaphase and anaphase are shown in which both daughter cells underwent subsequent mitosis (Fig 10A and C). Cells that did not divide and reverted are shown in Fig 10B and D. Daughter cells carrying the damaged chromatin took longer to divide than their counterparts without damaged chromatin (S3 B).

**Fig 9.**
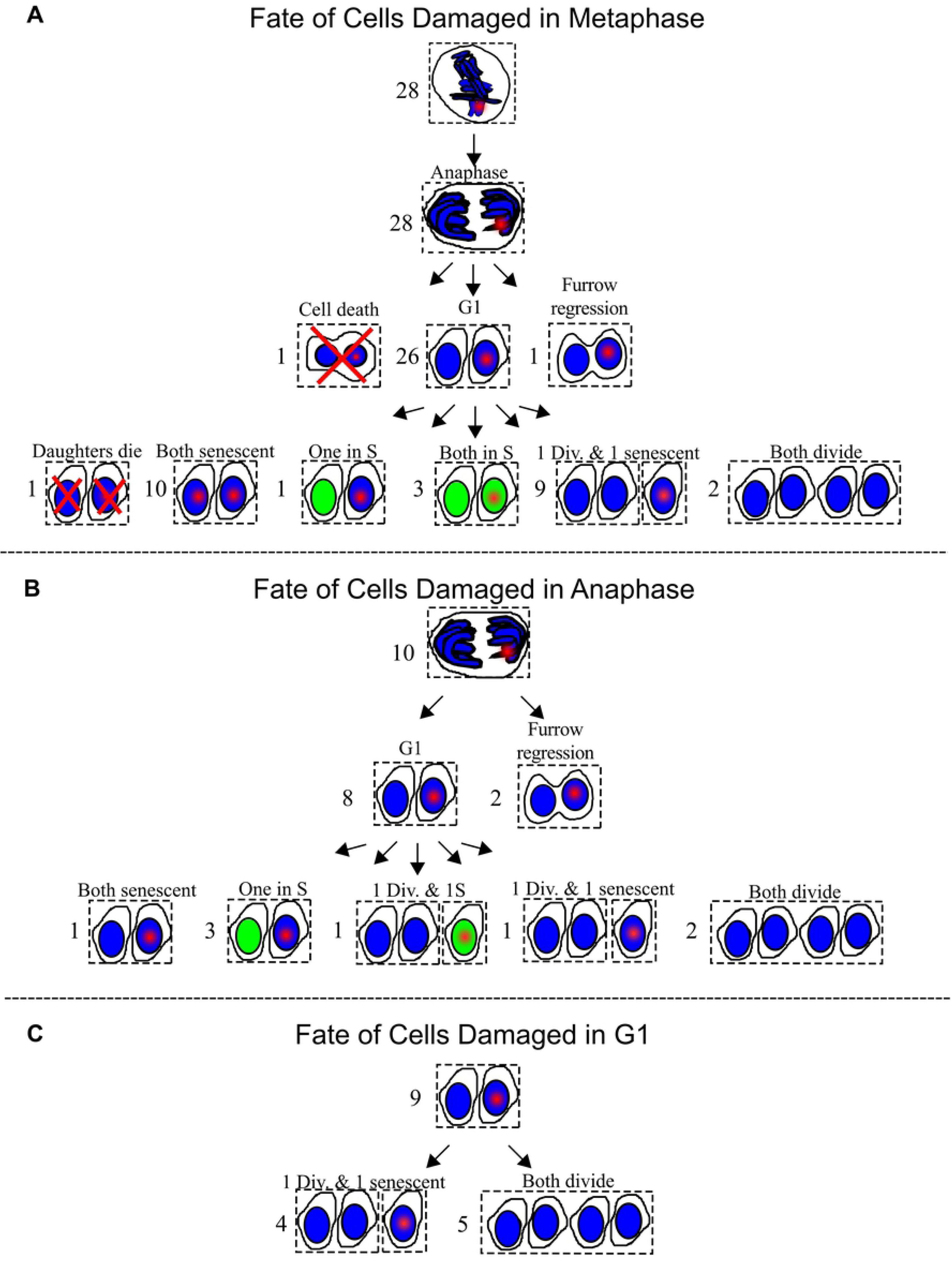
Fate of Cells Damaged in Mitosis and G1. (A) Twenty-eight metaphase cells were DNA damaged by the laser. A red point depicts laser damage. Twenty six out of the twenty-eight divided. One underwent furrow regression. Another underwent cleavage formation followed by cell death. The fates of daughters sets are shown below. A green nucleus marks S-phase. Divides is abbreviated as div. Six different outcomes are summarized: 1) both daughters die, 2) both daughters are senescent, 3) one daughter is in S and another is senescent, 4) both daughters are senescent, 5) one daughter divided and another is senescent, and 6) both daughters divide. (B) Ten cells were DNA damaged during Anaphase. Eight cells progressed into G1. Of the two that did not progress into G1, one underwent furrow regression, and another appears to have fused at a later point. Five outcomes for the daughters are summarized: 1) one set had both daughters in senescence, 2) three sets had one in S and the other in senescence, 3) one had one daughter divide and another in S phase, 4) another had one daughter divide and the other was senescent and 5) two sets had both daughters divide. (C) Nine G1 sisters were identified. One sister from each set was damaged. The cell containing the damaged nucleus has a red point. All of the undamaged sisters divided. Five of the damaged sisters divided.

**Fig 10.**
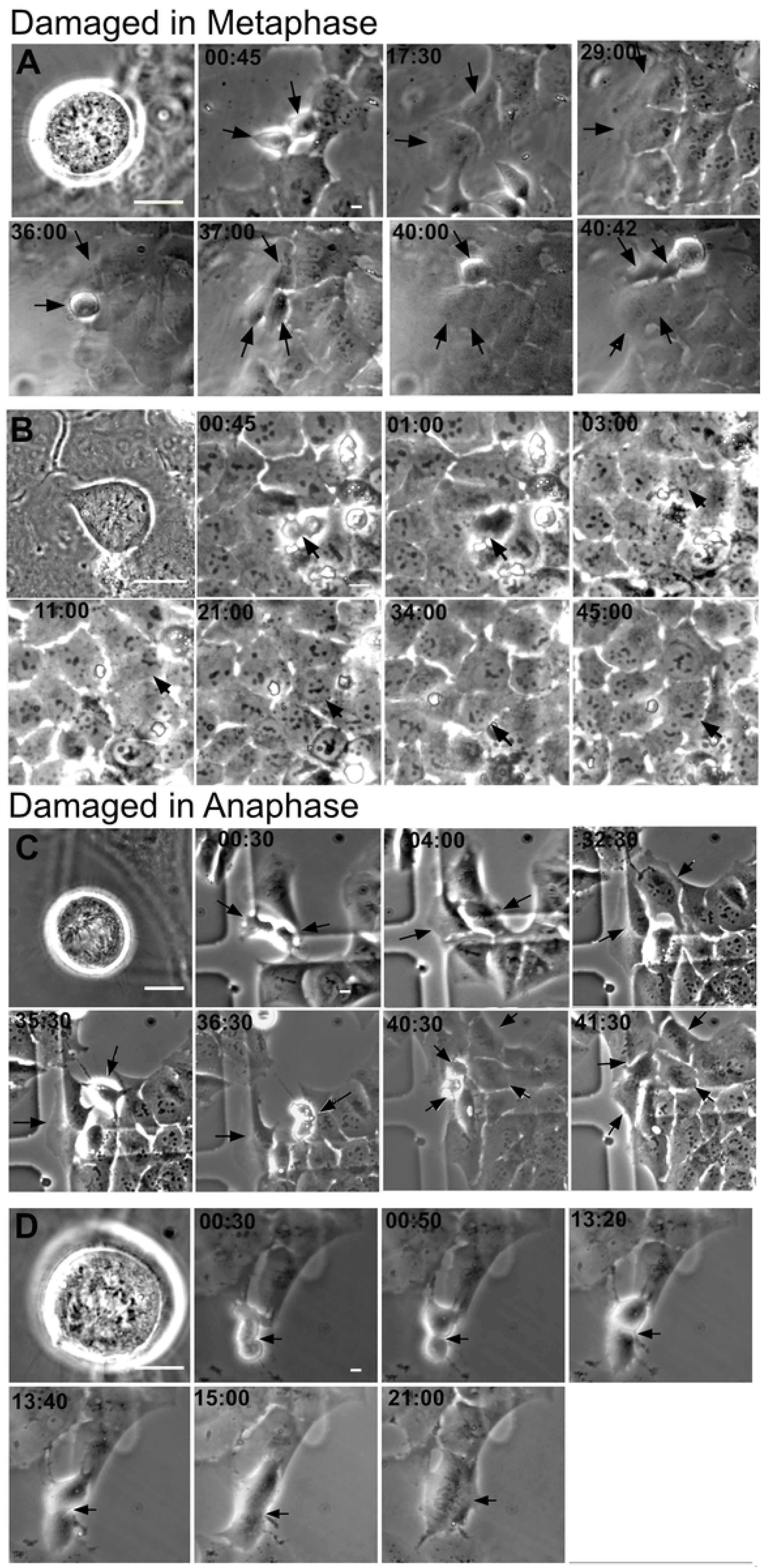
Time-lapse of cells damaged in mitosis. (A) Montage of a cell DNA damaged in metaphase whose daughters underwent mitosis at 36 and 40 hrs post division. Laser damage was created through a 63x objective. Therefore, cells appear larger on the first image. Subsequent images were taken with a 20x objective to broaden the field of view. (B) A metaphase DNA damaged cell whose furrow regressed at 1hr. (C) Montage of a cell damaged in Anaphase whose daughters divide at 36:30 and 40:30 hours. (D) An anaphase cell that appears to have divided, see 00:50 and 13:20. However, at 13:40 and 15:00 the cell begins to show furrow regression. Scale bar =10μm.

Damaged G1 cells were also followed to compare their ability to repair with that of mitotic cells. These cells were identified by following anaphase cells until completion of division and formation of two daughter cells. Of both daughter cells only one sister was damaged with the laser. However, both sisters were followed. Prior to fixation, all cells were incubated with EdU to check for S-phase status of cells that had not divided within the observation window. Fig 9 contains a summary of cell fates.

As expected all undamaged G1 sisters entered mitosis within the observation window. Out of the nine damaged cells, five divided i.e. 55% divided. Our results suggest that damage in metaphase is more deleterious than damage induced in anaphase or G1 (Fig 9C). Notwithstanding, these results demonstrate that a percentage (25%) of cells laser damaged in mitosis (metaphase and/or anaphase) are capable of undergoing a subsequent mitosis.

## Discussion

We present data demonstrating that strand breaks by the 780 nm NIR laser induce a DDR in mitotic cells and that various DNA repair proteins are recruited that are known to function in DSB repair during Interphase Fig 11. Additionally, we demonstrate that mitotic cells are capable of ongoing DNA repair synthesis when damage is elicited by an external source, laser or UV light. Furthermore, our results show that individual cells with damaged chromosomes are capable of progressing through the cell cycle and undergoing a subsequent cell division.

**Fig 11.**
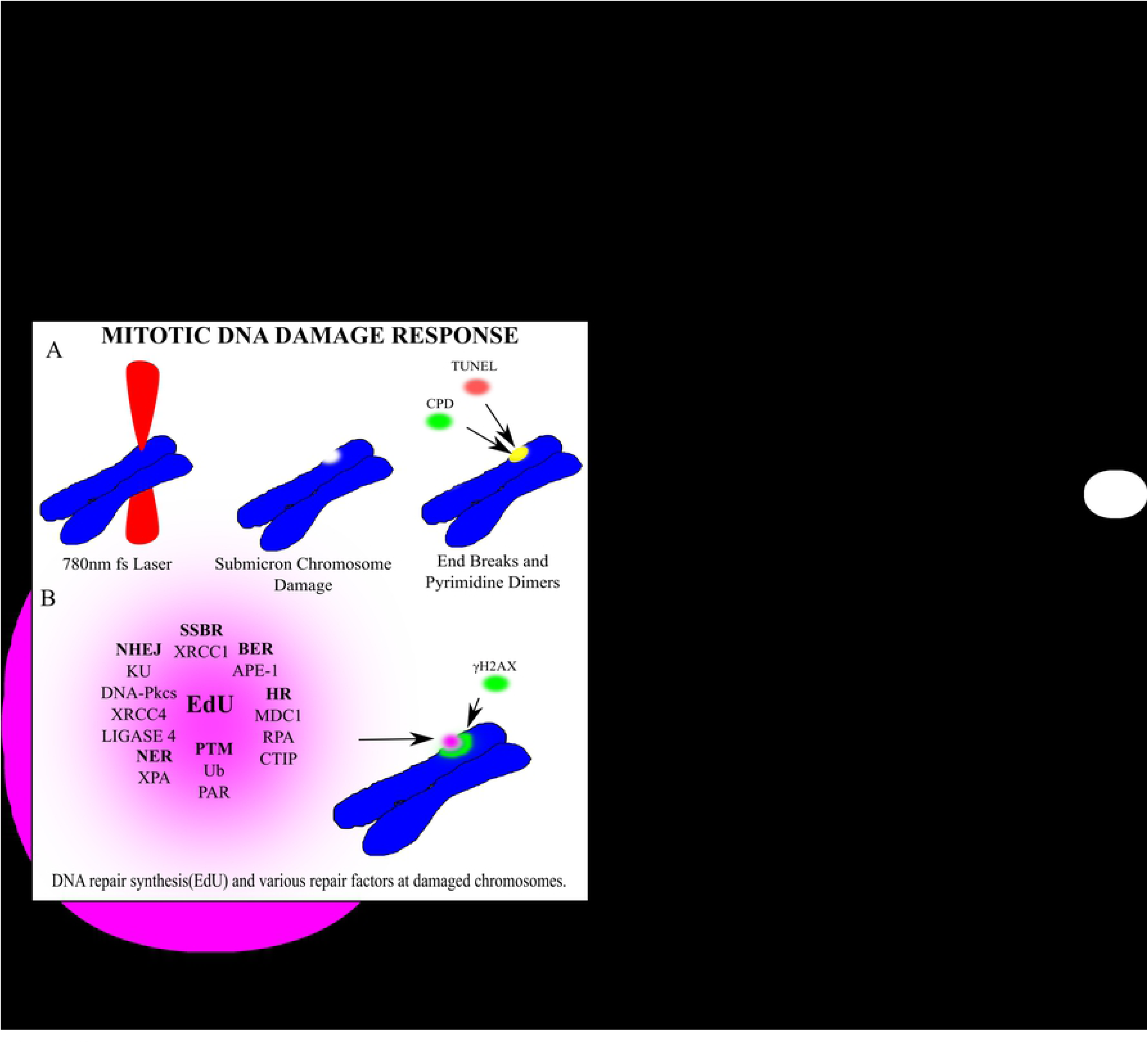
Article Summary: Mitotic DNA Damage Response. (A) A 780nm femtosecond laser was focused to a sub-micron region on a mitotic chromosome. End breaks detected via TUNEL assay and cyclo-butane pyrimidine dimers were found at the laser damage site. (B) Several factors cluster to the damage site. In bold are the repair pathway abbreviations that each factor is most closely associated with. Non-Homologous End Joining (NHEJ), Single Strand Break Repair (SSBR), Base Excision Repair(BER), Homologous Recombination(HR) Post translational modifications(PTM), Nucleotide excision repair(NER). DNA synthesis occurs at the damaged chromosome region as detected via EdU incorporation. Phosphorylated Histone γH2AX on Serine 139 marks double strand breaks and extends from the laser damage spot.

### The laser as a method to elucidate the DDR during mitosis

Using the highly focused NIR laser we have determined that the mitotic DNA damage response is more extensive than previously thought. This is likely due to a large amount of DNA damage in a small submicron volume, thus enabling detection of a high concentration of damage factors. Previous studies have utilized ionizing radiation and radiomimetic drugs to induce DSBs. Such studies did not show the accumulation of ubiquitin (Ub), RNF8, RNF168, BRCA1, 53BP1 to mitotic chromosomes [2, 5–14]. However, the results of those studies relied upon ionizing radiation induced foci formation which has been shown to be distinct from initial recruitment of the DNA damage-recognition factors and entails further clustering of proteins as well as amplification signals that surround damage sites [49, 66]. Nevertheless, we previously observed, using this same laser, that Ub was occurring at laser induced DSBs on mitotic chromosomes that coincided with KU staining [28, 38]. Additionally, the results in the present study show that BRCA1 and 53BP1 can localize to mitotic DNA damage sites. In line with this, Pederson et. Al. found 53BP1 foci on PICH associated ultrafine anaphase bridges and on chromosomes [67]. Therefore, the mechanisms regulating these interactions are more complex and likely depend on the type of damage, quantity and the phase in which damage is induced.

### Mitotic DDR may attempt to repair chromosomes by more than one pathway

Our results show that NHEJ was activated and was likely repairing some of the laser induced DNA damage. Interestingly, the KU signal was lower when the DNA ligase signal was greater. These results suggest NHEJ repair of DSBs. However, a decrease in KU binding may also be due to the abortive initiation of homologous recombination (HR). Mitotic cells may activate alternative-NHEJ, cells deficient in DNA-PKcs (a key component of NHEJ), still synthesized DNA. On the other hand, inhibition of PARP (a key player in alt-NHEJ) in these cells did not decrease DNA repair synthesis; instead, DNA repair synthesis appeared to increase. This result suggests that PARP has inhibitory effects on mitotic DNA repair and that HR may have been stimulated when both PARP and DNA-PKcs are compromised. Nevertheless, ATM inhibition alone did not decrease DNA repair synthesis. However, a decrease occurred when both ATM and DNA-PKcs activity was compromised. This is not surprising given that ATM and DNA PKcs are key to the DSB repair response in that they can phosphorylate γH2AX[68]. Further, ATM activity was recently shown to be important for NER [69]. Therefore, DNA repair synthesis may occur in response to different pathways depending on the cell conditions and/or availability of specific repair proteins. Future studies are required to investigate the ability of mitotic cells to undergo NER. Our results showing that the laser is capable of inducing UV type damage demonstrate this to be an excellent tool toward that end.

Further, the complex nature of laser damage likely leads to the activity of multiple repair pathways. Consistent with this, factors involved in BER/SSB repair APE1 and XRCC1 were also detected at mitotic chromosomes after laser damage. These results are consistent with a previous study that found similar results when mitotic cells were treated with hydrogen peroxide to induce DNA damage [70].

Though HR repair in mitosis is repressed and sister chromatids are separately condensed, the chromatin may be prepared for HR repair by end resection visualized through increased RPA binding and expansion. Even when cells are kept in mitosis for prolonged periods by nocodazole, downstream repair activation by RAD51 filament formation is not observed; nor is it observed in late stages of mitosis such as telophase. However, consistent with previous non-laser studies, RAD51 is more likely to be recruited to mitotic DNA damage when CDK1 is inhibited. Greater RAD51 recruitment was readily observed in the absence of DNA PKcs. These results demonstrate that DNA-PKcs is critical to blocking RAD51 recruitment during mitosis and that laser damaged mitotic chromosomes may have a greater ability to recruit NHEJ factors.

### Mitotic DNA repair synthesis via different means

Mitotic DNA repair synthesis is not a laser specific phenomenon. Treatment of mitotic cells with a conventional UV light source also resulted in EdU incorporation, indicating that this type of DNA repair synthesis is not a laser damage-specific phenomenon. Previous studies have shown that replication stress induced DNA damage can lead to DNA repair synthesis in very early prophase but not later phases of mitosis. Cells synchronized in nocodazole did not undergo DNA repair synthesis [16, 17]. However, the mechanism of DNA repair synthesis observed in our studies likely differs from those studies in that (1) our damage is inflicted in later prophase, metaphase and anaphase and (2) our damage is not due to active replication stress induced repair. The process described in Bhowmick et al., 2016 and Minocherhomiji et al., 2015 is dependent on Rad52 and MUS81-EME1. It may be interesting to determine whether DNA damage induced in early prophase by means other than replication stress requires the same factors, or determine how inhibition of ATM and DNA-PKcs affects replication stress induced repair that is observed in early prophase.

### Damage induced in Metaphase is more deleterious for cell division

We investigated the ability of cells damaged in mitosis to enter a second mitosis. This is an important question raised by early studies using a different laser (argon ion laser emitting 488 or 514 nm light) where mitotic cells with focal damage to a single chromosome were still able to undergo a subsequent mitosis and produce apparently normal cells [26].This is significant because entry into second mitosis is indicative of checkpoint recovery and thus DNA repair to the point that it is no longer halting cell cycle progression. Under our laser conditions, we observed that a significant percentage (25%) of cells damaged in mitosis undergo a second division. This fraction varies depending on the phase in which damage was induced. Cells damaged in metaphase were less likely to enter a second division in comparison to cells damaged in anaphase or G1. This may be caused by irradiation of a more compacted metaphase chromosome resulting in more DNA damage than in G1. The DNA repair pathway choice in mitosis can be another important factor.

In conclusion, our results show that (1) mitotic cells are capable of DNA repair synthesis, (2) factors from multiple repair pathways can localize to mitotic DNA damage; therefore, DNA repair synthesis may be a result of one or more repair mechanisms, (3) DNA repair synthesis is compromised when ATM and DNA-Pkcs are inhibited; individual inhibition did not show significant changes, and (4) 25% of cells carrying mitotic DNA damage are able to undergo DNA repair, progress through the cell cycle, and enter a subsequent division.

## Supplemental Information

S1 Assessment of the mitotic DNA damage response

S2 DNA damage response in different cell lines (M059K, M059J and CFPAC1).

S3 Time to cell division summary and antibody list

## Acknowledgements

A special thanks to Arthur Forer, PhD (York University, Toronto, ON) and Phang- Lang Chen (UC-Irvine, Irvine, Ca) for thought provoking conversations on mitotic responses to damage.

## Abbreviations

DSB: double strand break
DDR: DNA damage response
HR: Homologous Recombination Repair
NHEJ: Non-homologous end joining
NER: Nucleotide Excision Repair
EdU: 5-Ethynyl-2’-deoxyuridine
DNA PKcsi: DNA PKcs Inhibitor
ATMi: ATM inhibitor
PARPi: PARP Inhibitor

## References

1. Castedo M, Perfettini JL, Roumier T, Valent A, Raslova H, Yakushijin K, et al. Mitotic catastrophe constitutes a special case of apoptosis whose suppression entails aneuploidy. Oncogene. 2004;23(25):4362–70. doi: 10.1038/sj.onc.1207572. PubMed PMID: 15048075.

2. Orthwein A, Fradet-Turcotte A, Noordermeer SM, Canny MD, Brun CM, Strecker J, et al. Mitosis inhibits DNA double-strand break repair to guard against telomere fusions. Science. 2014;344(6180):189–93. doi: 10.1126/science.1248024. PubMed PMID: 24652939.

3. Crasta K, Ganem NJ, Dagher R, Lantermann AB, Ivanova EV, Pan Y, et al. DNA breaks and chromosome pulverization from errors in mitosis. Nature. 2012;482(7383):53-8. doi: 10.1038/nature10802. PubMed PMID: 22258507; PubMed Central PMCID: PMCPMC3271137.

4. Bakhoum SF, Kabeche L, Compton DA, Powell SN, Bastians H. Mitotic DNA Damage Response: At the Crossroads of Structural and Numerical Cancer Chromosome Instabilities. Trends Cancer. 2017;3(3):225–34. doi: 10.1016/j.trecan.2017.02.001. PubMed PMID: 28718433; PubMed Central PMCID: PMCPMC5518619.

5. Giunta S, Belotserkovskaya R, Jackson SP. DNA damage signaling in response to double-strand breaks during mitosis. J Cell Biol. 2010;190(2):197–207. Epub 2010/07/28. doi: jcb.200911156 [pii] 10.1083/jcb.200911156. PubMed PMID: 20660628.

6. Huang X, Tran T, Zhang L, Hatcher R, Zhang P. DNA damage-induced mitotic catastrophe is mediated by the Chk1-dependent mitotic exit DNA damage checkpoint. Proceedings of the National Academy of Sciences of the United States of America. 2005;102(4):1065–70. Epub 2005/01/15. doi: 10.1073/pnas.0409130102. PubMed PMID: 15650047; PubMed Central PMCID: PMC545827.

7. Lee DH, Acharya SS, Kwon M, Drane P, Guan Y, Adelmant G, et al. Dephosphorylation enables the recruitment of 53BP1 to double-strand DNA breaks. Mol Cell. 2014;54(3):512–25. doi: 10.1016/j.molcel.2014.03.020. PubMed PMID: 24703952; PubMed Central PMCID: PMCPMC4030556.

8. Nelson G, Buhmann M, von Zglinicki T. DNA damage foci in mitosis are devoid of 53BP1. Cell Cycle. 2009;8(20):3379–83. Epub 2009/10/07. PubMed PMID: 19806024.

9. Wei Zhang GP, Shiaw-Yin Lin and Pumin Zhang. DNA damage response is suppressed by high Cdk1 activity in mitotic mammalian cells. Journal of Biological Chemistry. 2011;107(15):6870–5. Epub March, 29, 2010.

10. Zhang W, Peng G, Lin SY, Zhang P. DNA Damage Response Is Suppressed by the High Cyclin-dependent Kinase 1 Activity in Mitotic Mammalian Cells. Journal of Biological Chemistry. 2011;286(41):35899–905. doi: 10.1074/jbc.M111.267690.

11. van Vugt MA, Gardino AK, Linding R, Ostheimer GJ, Reinhardt HC, Ong SE, et al. A mitotic phosphorylation feedback network connects Cdk1, Plk1, 53BP1, and Chk2 to inactivate the G(2)/M DNA damage checkpoint. PLoS Biol. 2010;8(1):e1000287. doi: 10.1371/journal.pbio.1000287. PubMed PMID: 20126263; PubMed Central PMCID: PMCPMC2811157.

12. Benada J, Burdova K, Lidak T, von Morgen P, Macurek L. Polo-like kinase 1 inhibits DNA damage response during mitosis. Cell Cycle. 2015;14(2):219–31. doi: 10.4161/15384101.2014.977067. PubMed PMID: 25607646; PubMed Central PMCID: PMCPMC4613155.

13. Peterson SE, Li Y, Chait BT, Gottesman ME, Baer R, Gautier J. Cdk1 uncouples CtIP-dependent resection and Rad51 filament formation during M- phase double-strand break repair. The Journal of Cell Biology. 2011;194(5):705–20. doi: 10.1083/jcb.201103103.

14. Terasawa M, Shinohara A, Shinohara M. Canonical non-homologous end joining in mitosis induces genome instability and is suppressed by M-phase- specific phosphorylation of XRCC4. PLoS Genet. 2014;10(8):e1004563. doi: 10.1371/journal.pgen.1004563. PubMed PMID: 25166505; PubMed Central PMCID: PMCPMC4148217.

15. Feng W, Jasin M. Homologous Recombination and Replication Fork Protection: BRCA2 and More! Cold Spring Harb Symp Quant Biol. 2017;82:329–38. doi: 10.1101/sqb.2017.82.035006. PubMed PMID: 29686033; PubMed Central PMCID: PMCPMC6333483.

16. Bhowmick R, Minocherhomji S, Hickson ID. RAD52 Facilitates Mitotic DNA Synthesis Following Replication Stress. Mol Cell. 2016;64(6):1117–26. doi: 10.1016/j.molcel.2016.10.037. PubMed PMID: 27984745.

17. Minocherhomji S, Ying S, Bjerregaard VA, Bursomanno S, Aleliunaite A, Wu W, et al. Replication stress activates DNA repair synthesis in mitosis. Nature. 2015;528(7581):286–90. doi: 10.1038/nature16139. PubMed PMID: 26633632.

18. Ying S, Minocherhomji S, Chan KL, Palmai-Pallag T, Chu WK, Wass T, et al. MUS81 promotes common fragile site expression. Nat Cell Biol. 2013;15(8):1001–7. doi: 10.1038/ncb2773. PubMed PMID: 23811685.

19. Hanada K, Budzowska M, Davies SL, van Drunen E, Onizawa H, Beverloo HB, et al. The structure-specific endonuclease Mus81 contributes to replication restart by generating double-strand DNA breaks. Nat Struct Mol Biol. 2007;14(11):1096–104. doi: 10.1038/nsmb1313. PubMed PMID: 17934473.

20. Davies SL, North PS, Hickson ID. Role for BLM in replication-fork restart and suppression of origin firing after replicative stress. Nat Struct Mol Biol. 2007;14(7):677–9. doi: 10.1038/nsmb1267. PubMed PMID: 17603497.

21. Min J, Wright WE, Shay JW. Alternative Lengthening of Telomeres Mediated by Mitotic DNA Synthesis Engages Break-Induced Replication Processes. Mol Cell Biol. 2017;37(20). doi: 10.1128/MCB.00226-17. PubMed PMID: 28760773; PubMed Central PMCID: PMCPMC5615184.

22. Feng W, Jasin M. BRCA2 suppresses replication stress-induced mitotic and G1 abnormalities through homologous recombination. Nat Commun. 2017;8(1):525. doi: 10.1038/s41467-017-00634-0. PubMed PMID: 28904335; PubMed Central PMCID: PMCPMC5597640.

23. Berns MW. Directed chromosome loss by laser microirradiation. Science. 1974;186(4165):700–5. Epub 1974/11/22. PubMed PMID: 4607753.

24. Berns MW. Laser microirradiation of chromosomes. Cold Spring Harb Symp Quant Biol. 1974;38:165–74. Epub 1974/01/01. PubMed PMID: 4133983.

25. Berns MW. The laser microbeam as a probe for chromatin structure and function. Methods Cell Biol. 1978;18:277–94. Epub 1978/01/01. PubMed PMID: 683019.

26. Berns MW, Cheng WK, Hoover G. Cell division after laser microirradiation of mitotic chromosomes. Nature. 1971;233(5315):122–3. Epub 1971/09/10. PubMed PMID: 12058751.

27. Berns MW, Olson RS, Rounds DE. In vitro production of chromosomal lesions with an argon laser microbeam. Nature. 1969;221(5175):74–5. Epub 1969/01/04. PubMed PMID: 4882051.

28. Gomez-Godinez V, Wu T, Sherman AJ, Lee CS, Liaw LH, Zhongsheng Y, et al. Analysis of DNA double-strand break response and chromatin structure in mitosis using laser microirradiation. Nucleic Acids Res. 38(22):e202. Epub 2010/10/07. doi: gkq836 [pii] 10.1093/nar/gkq836. PubMed PMID: 20923785; PubMed Central PMCID: PMC3001094.

29. Silva BA, Stambaugh JR, Yokomori K, Shah JV, Berns MW. DNA damage to a single chromosome end delays anaphase onset. J Biol Chem. 2014;289(33):22771–84. doi: 10.1074/jbc.M113.535955. PubMed PMID: 24982423; PubMed Central PMCID: PMCPMC4132783.

30. Berns MW, Chong LK, Hammer-Wilson M, Miller K, Siemens A. Genetic microsurgery by laser: establishment of a clonal population of rat kangaroo cells (PTK2) with a directed deficiency in a chromosomal nucleolar organizer. Chromosoma. 1979;73(1):1–8. Epub 1979/06/21. PubMed PMID: 487905.

31. Kong X, Mohanty SK, Stephens J, Heale JT, Gomez-Godinez V, Shi LZ, et al. Comparative analysis of different laser systems to study cellular responses to DNA damage in mammalian cells. Nucleic Acids Res. 2009;37(9):e68. doi: 10.1093/nar/gkp221. PubMed PMID: 19357094; PubMed Central PMCID: PMCPMC2685111.

32. Saquilabon Cruz GM, Kong X, Silva BA, Khatibzadeh N, Thai R, Berns MW, et al. Femtosecond near-infrared laser microirradiation reveals a crucial role for PARP signaling on factor assemblies at DNA damage sites. Nucleic Acids Res. 2016;44(3):e27. doi: 10.1093/nar/gkv976. PubMed PMID: 26424850; PubMed Central PMCID: PMCPMC4756852.

33. Gomez-Godinez V, Wakida NM, Dvornikov AS, Yokomori K, Berns MW. Recruitment of DNA damage recognition and repair pathway proteins following near-IR femtosecond laser irradiation of cells. J Biomed Opt. 2007;12(2):020505. doi: 10.1117/1.2717684. PubMed PMID: 17477704.

34. Gomez-Godinez V, Wu T, Sherman AJ, Lee CS, Liaw LH, Zhongsheng Y, et al. Analysis of DNA double-strand break response and chromatin structure in mitosis using laser microirradiation. Nucleic Acids Res. 2010;38(22):e202. doi: 10.1093/nar/gkq836. PubMed PMID: 20923785; PubMed Central PMCID: PMCPMC3001094.

35. Kong X, Ball AR, Jr., Pham HX, Zeng W, Chen HY, Schmiesing JA, et al. Distinct functions of human cohesin-SA1 and cohesin-SA2 in double-strand break repair. Mol Cell Biol. 2014;34(4):685–98. doi: 10.1128/MCB.01503-13. PubMed PMID: 24324008; PubMed Central PMCID: PMCPMC3911484.

36. Kong X, Stephens J, Ball AR, Jr., Heale JT, Newkirk DA, Berns MW, et al. Condensin I recruitment to base damage-enriched DNA lesions is modulated by PARP1. PLoS One. 2011;6(8):e23548. doi: 10.1371/journal.pone.0023548. PubMed PMID: 21858164; PubMed Central PMCID: PMCPMC3155556.

37. Silva BA, Stambaugh JR, Berns MW. Targeting telomere-containing chromosome ends with a near-infrared femtosecond laser to study the activation of the DNA damage response and DNA damage repair pathways. J Biomed Opt. 2013;18(9):095003. doi: 10.1117/1.JBO.18.9.095003. PubMed PMID: 24064949; PubMed Central PMCID: PMCPMC3782557.

38. Mari PO, Florea BI, Persengiev SP, Verkaik NS, Bruggenwirth HT, Modesti M, et al. Dynamic assembly of end-joining complexes requires interaction between Ku70/80 and XRCC4. Proc Natl Acad Sci U S A. 2006;103(49):18597–602. Epub 2006/11/25. doi: 0609061103 [pii] 10.1073/pnas.0609061103. PubMed PMID: 17124166; PubMed Central PMCID: PMC1693708.

39. Saquilabon Cruz GM, Kong X, Silva BA, Khatibzadeh N, Thai R, Berns MW, et al. Femtosecond near-infrared laser microirradiation reveals a crucial role for PARP signaling on factor assemblies at DNA damage sites. Nucleic Acids Res. 2015. doi: 10.1093/nar/gkv976. PubMed PMID: 26424850.

40. Wakida NM, Lee CS, Botvinick ET, Shi LZ, Dvornikov A, Berns MW. Laser nanosurgery of single microtubules reveals location-dependent depolymerization rates. Journal of biomedical optics. 2007;12(2):024022. Epub 2007/05/05. doi: 10.1117/1.2718920. PubMed PMID: 17477737.

41. Gomez-Godinez V, Wakida NM, Dvornikov AS, Yokomori K, Berns MW. Recruitment of DNA damage recognition and repair pathway proteins following near-IR femtosecond laser irradiation of cells. Journal of biomedical optics. 2007;12(2):020505. Epub 2007/05/05. doi: 10.1117/1.2717684. PubMed PMID: 17477704.

42. Botvinick EL, Venugopalan V, Shah JV, Liaw LH, Berns MW. Controlled ablation of microtubules using a picosecond laser. Biophys J. 2004;87(6):4203–12. Epub 2004/09/30. doi: 10.1529/biophysj.104.049528 S0006-3495(04)73884-9 [pii]. PubMed PMID: 15454403; PubMed Central PMCID: PMC1304929.

43. Rasband W.S. Image J Bethesda, Maryland, USA: US. National Institutes of Health; 1997-2009. Available from: http://rsb.info.nih.gov/ij/,.

44. London RE. The structural basis of XRCC1-mediated DNA repair. DNA Repair (Amst). 2015;30:90–103. doi: 10.1016/j.dnarep.2015.02.005. PubMed PMID: 25795425; PubMed Central PMCID: PMCPMC5580684.

45. Rogakou EP, Pilch DR, Orr AH, Ivanova VS, Bonner WM. DNA double- stranded breaks induce histone H2AX phosphorylation on serine 139. J Biol Chem. 1998;273(10):5858–68. Epub 1998/04/16. PubMed PMID: 9488723.

46. Rogakou EP, Boon C, Redon C, Bonner WM. Megabase chromatin domains involved in DNA double-strand breaks in vivo. J Cell Biol. 1999;146(5):905–16. Epub 1999/09/09. PubMed PMID: 10477747; PubMed Central PMCID: PMC2169482.

47. Iarovaia OV, Rubtsov M, Ioudinkova E, Tsfasman T, Razin SV, Vassetzky YS. Dynamics of double strand breaks and chromosomal translocations. Mol Cancer. 2014;13:249. doi: 10.1186/1476-4598-13-249. PubMed PMID: 25404525; PubMed Central PMCID: PMCPMC4289179.

48. Uziel T, Lerenthal Y, Moyal L, Andegeko Y, Mittelman L, Shiloh Y. Requirement of the MRN complex for ATM activation by DNA damage. EMBO J. 2003;22(20):5612–21. Epub 2003/10/09. doi: 10.1093/emboj/cdg541. PubMed PMID: 14532133; PubMed Central PMCID: PMC213795.

49. Kim JS, Krasieva TB, Kurumizaka H, Chen DJ, Taylor AM, Yokomori K. Independent and sequential recruitment of NHEJ and HR factors to DNA damage sites in mammalian cells. J Cell Biol. 2005;170(3):341–7. Epub 2005/08/03. doi: jcb.200411083 [pii] 10.1083/jcb.200411083. PubMed PMID: 16061690; PubMed Central PMCID: PMC2171485.

50. Lukas C, Melander F, Stucki M, Falck J, Bekker-Jensen S, Goldberg M, et al. Mdc1 couples DNA double-strand break recognition by Nbs1 with its H2AX- dependent chromatin retention. Embo J. 2004;23(13):2674–83. PubMed PMID: 15201865.

51. Giunta S, Belotserkovskaya R, Jackson SP. DNA damage signaling in response to double-strand breaks during mitosis. J Cell Biol. 190(2):197–207. Epub 2010/07/28. doi: jcb.200911156 [pii] 10.1083/jcb.200911156. PubMed PMID: 20660628.

52. Stewart GS. Solving the RIDDLE of 53BP1 recruitment to sites of damage. Cell Cycle. 2009;8(10):1532–8. Epub 2009/04/18. doi: 8351 [pii]. PubMed PMID: 19372751.

53. Stewart GS, Panier S, Townsend K, Al-Hakim AK, Kolas NK, Miller ES, et al. The RIDDLE syndrome protein mediates a ubiquitin-dependent signaling cascade at sites of DNA damage. Cell. 2009;136(3):420–34. Epub 2009/02/11. doi: S0092-8674(09)00005-1 [pii] 10.1016/j.cell.2008.12.042. PubMed PMID: 19203578.

54. Perrault R, Wang H, Wang M, Rosidi B, Iliakis G. Backup pathways of NHEJ are suppressed by DNA-PK. Journal of cellular biochemistry. 2004;92(4):781–94. Epub 2004/06/24. doi: 10.1002/jcb.20104. PubMed PMID: 15211575.

55. Mansour WY, Rhein T, Dahm-Daphi J. The alternative end-joining pathway for repair of DNA double-strand breaks requires PARP1 but is not dependent upon microhomologies. Nucleic Acids Res. 2010;38(18):6065–77. doi: 10.1093/nar/gkq387. PubMed PMID: 20483915; PubMed Central PMCID: PMCPMC2952854.

56. Wang M, Wu W, Wu W, Rosidi B, Zhang L, Wang H, et al. PARP-1 and Ku compete for repair of DNA double strand breaks by distinct NHEJ pathways. Nucleic Acids Res. 2006;34(21):6170–82. doi: 10.1093/nar/gkl840. PubMed PMID: 17088286; PubMed Central PMCID: PMCPMC1693894.

57. Saberi A, Hochegger H, Szuts D, Lan L, Yasui A, Sale JE, et al. RAD18 and poly(ADP-ribose) polymerase independently suppress the access of nonhomologous end joining to double-strand breaks and facilitate homologous recombination-mediated repair. Mol Cell Biol. 2007;27(7):2562–71. doi: 10.1128/MCB.01243-06. PubMed PMID: 17242200; PubMed Central PMCID: PMCPMC1899888.

58. Dominguez-Bendala J, Masutani M, McWhir J. Down-regulation of PARP- 1, but not of Ku80 or DNA-PKcs’, results in higher gene targeting efficiency. Cell Biol Int. 2006;30(4):389–93. doi: 10.1016/j.cellbi.2005.12.005. PubMed PMID: 16504547.

59. Bakr A, Oing C, Kocher S, Borgmann K, Dornreiter I, Petersen C, et al. Involvement of ATM in homologous recombination after end resection and RAD51 nucleofilament formation. Nucleic Acids Res. 2015;43(6):3154–66. doi: 10.1093/nar/gkv160. PubMed PMID: 25753674; PubMed Central PMCID: PMCPMC4381069.

60. Beucher A, Birraux J, Tchouandong L, Barton O, Shibata A, Conrad S, et al. ATM and Artemis promote homologous recombination of radiation-induced DNA double-strand breaks in G2. EMBO J. 2009;28(21):3413–27. doi: 10.1038/emboj.2009.276. PubMed PMID: 19779458; PubMed Central PMCID: PMCPMC2752027.

61. Moynahan ME, Jasin M. Mitotic homologous recombination maintains genomic stability and suppresses tumorigenesis. Nature reviews Molecular cell biology. 2010;11(3):196–207. Epub 2010/02/24. doi: 10.1038/nrm2851. PubMed PMID: 20177395; PubMed Central PMCID: PMC3261768.

62. Peterson SE, Li Y, Chait BT, Gottesman ME, Baer R, Gautier J. Cdk1 uncouples CtIP-dependent resection and Rad51 filament formation during M- phase double-strand break repair. J Cell Biol. 194(5):705–20. Epub 2011/09/07. doi: jcb.201103103 [pii] 10.1083/jcb.201103103. PubMed PMID: 21893598; PubMed Central PMCID: PMC3171114.

63. Vassilev LT. Selective small-molecule inhibitor reveals critical mitotic functions of human CDK1. Proceedings of the National Academy of Sciences. 2006;103(28):10660–5. doi: 10.1073/pnas.0600447103.

64. Potts PR, Porteus MH, Yu H. Human SMC5/6 complex promotes sister chromatid homologous recombination by recruiting the SMC1/3 cohesin complex to double-strand breaks. EMBO J. 2006;25(14):3377–88. doi: 10.1038/sj.emboj.7601218. PubMed PMID: 16810316; PubMed Central PMCID: PMCPMC1523187.

65. Nishiyama T, Sykora MM, Huis in ’t Veld PJ, Mechtler K, Peters JM. Aurora B and Cdk1 mediate Wapl activation and release of acetylated cohesin from chromosomes by phosphorylating Sororin. Proc Natl Acad Sci U S A. 2013;110(33):13404–9. doi: 10.1073/pnas.1305020110. PubMed PMID: 23901111; PubMed Central PMCID: PMCPMC3746921.

66. Celeste A, Fernandez-Capetillo O, Kruhlak MJ, Pilch DR, Staudt DW, Lee A, et al. Histone H2AX phosphorylation is dispensable for the initial recognition of DNA breaks. Nat Cell Biol. 2003;5(7):675–9. Epub 2003/06/07. doi: 10.1038/ncb1004ncb1004 [pii]. PubMed PMID: 12792649.

67. Pedersen RT, Kruse T, Nilsson J, Oestergaard VH, Lisby M. TopBP1 is required at mitosis to reduce transmission of DNA damage to G1 daughter cells. J Cell Biol. 2015;210(4):565–82. doi: 10.1083/jcb.201502107. PubMed PMID: 26283799; PubMed Central PMCID: PMCPMC4539992.

68. Stiff T, O’Driscoll M, Rief N, Iwabuchi K, Lobrich M, Jeggo PA. ATM and DNA-PK function redundantly to phosphorylate H2AX after exposure to ionizing radiation. Cancer Res. 2004;64(7):2390–6. PubMed PMID: 15059890.

69. Wakasugi M, Sasaki T, Matsumoto M, Nagaoka M, Inoue K, Inobe M, et al. Nucleotide excision repair-dependent DNA double-strand break formation and ATM signaling activation in mammalian quiescent cells. J Biol Chem. 2014;289(41):28730–7. doi: 10.1074/jbc.M114.589747. PubMed PMID: 25164823; PubMed Central PMCID: PMCPMC4192521.

70. Okano S, Lan L, Tomkinson AE, Yasui A. Translocation of XRCC1 and DNA ligase IIIalpha from centrosomes to chromosomes in response to DNA damage in mitotic human cells. Nucleic acids research. 2005;33(1):422–9. Epub 2005/01/18. doi: 10.1093/nar/gki190. PubMed PMID: 15653642; PubMed Central PMCID: PMC546168.

